# Genomics-driven monitoring of *Fraxinus latifolia* (Oregon Ash) for conservation and EAB-resistance breeding

**DOI:** 10.1101/2024.08.22.609160

**Authors:** Anthony E. Melton, Trevor M. Faske, Richard A. Sniezko, Tim Thibault, Wyatt Williams, Thomas Parchman, Jill A. Hamilton

## Abstract

Understanding the evolutionary processes underlying range-wide genomic variation is critical to designing effective conservation and restoration strategies. Evaluating the influence of connectivity, demographic change, and environmental adaptation for threatened species can be invaluable to proactive conservation of evolutionary potential. In this study, we assessed genomic variation across the range of *Fraxinus latifolia*, a foundational riparian tree native to western North America recently exposed to the invasive emerald ash borer (*Agrilus planipennis*; EAB). Over 1,000 individuals from 61 populations were sequenced using reduced representation (ddRAD-seq) across the species’ range. Strong population structure was evident along a latitudinal gradient, with population connectivity largely maintained along central valley river systems, and a center of diversity coinciding with major river systems central to the species’ range. Despite evidence of connectivity, estimates of nucleotide diversity and effective population size were low across all populations, suggesting the patchy distribution of *F. latifolia* populations may impact its long-term evolutionary potential. Range-wide estimates of genomic offset, which indicate genomic change required to adjust to future climate projections, were greatest in the eastern and lowest in the southern portions of the species’ range, suggesting the regional distribution of genomic variation may impact evolutionary potential longer-term. To preserve evolutionary capacity across populations needed for development of climate-resilient, EAB-resistant breeding programs, prioritizing conservation of range-wide genomic diversity will provide a foundation for species management long-term.

## Introduction

The International Union for Conservation of Nature (IUCN) recently identified the maintenance of intraspecific genomic variation for species of conservation concern as a “hidden biodiversity crisis” (reviewed in Des Roches et al., 2021; Hoban et al., 2021). As the raw material upon which natural selection acts, maintenance of genomic variation is critical to maintaining evolutionary potential (reviewed in Kardos et al., 2021; McLaughlin et al., 2024). The combination of local extinctions, demographic declines, and selection in response to anthropogenic influences are threatening the persistence of intraspecific population genomic variation globally (Des Roches et al., 2021). Thus, to preserve long-term evolutionary potential for species at risk, understanding the evolutionary processes contributing to standing genomic variation is necessary. Landscape genomic analyses can inform conservation and restoration by assessing species-wide diversity and differentiation (Laporte & Charlesworth, 2002; Vandergast et al., 2006; Nock et al., 2011; Li et al., 2019; Osuna-Mascaró et al., 2023; reviewed in Savolainen et al., 2007; Hoban et al., 2016), estimating contemporary and predicting genotype-environment relationships (Meek et al., 2022; St. Clair et al., 2022), and evaluating effective population size (reviewed in Waples, 2010) and relatedness (Félix-Valdez et al., 2016; Tabas Madrid et al., 2018; Ruiz Mondragon et al., 2023), needed to inform conservation and restoration strategies. Such analyses can be invaluable to informing proactive conservation strategies, including the expansion of *ex situ* collections and establishment of genecological resources that can be foundational to restoring species at risk (Cavender et al., 2015; Diaz-Martin et al., 2023; reviewed in VanWallendael et al., 2022).

As a keystone tree species, ash (*Fraxinus* L.; Oleaceae) species that provide essential ecosystem services; including carbon sequestration, nutrient, and water cycling across North America forests (Barstow et al., 2018; Gandhi & Herms, 2010; Hausman et al., 2010; Herms & McCullough, 2014). However, the introduction of the emerald ash borer (EAB; *Agrilus planipennis* Fairmaire; Buprestidae) and its associated impacts have decimated *Fraxinus* species across much of eastern North America, causing one of the costliest forest insect invasions to date across the United States (Kovacs et al., 2010; Semizer-Cuming et al., 2019). With the continued range expansion of EAB, officially observed in Oregon, USA in 2022 and British Columbia, Canada in 2024, efforts to conserve and restore *Fraxinus* species have emphasized *ex situ* collections to preserve contemporary standing genomic variation and germplasm maintenance to lay a foundation for EAB-resistance breeding and future restoration (Carlson et al., 2014; Hamilton et al., 2020). Thus, genomics-driven monitoring of extant populations of *Fraxinus latifolia* Benth. (Oregon ash) may be leveraged to understand the demographic history of the species and how different neutral and non-neutral evolutionary processes have shaped standing genomic variation (Di Santo et al., 2022; Bolte et al., 2024). In addition, genotype-environment associations (GEA) enable characterization of the role of environmental adaptation in shaping contemporary genomic structure (reviewed in Wang & Bradburd, 2014; Lasky et al. 2023).

Modeling the relationship between genotypic and environmental variation provides the opportunity to evaluate the impact environmental change may have to genotype-environment associations, quantifying potential genomic mismatches under future climate conditions (i.e., genomic offset; Jia et al., 2020; Borrell et al., 2020; Mahony et al., 2020; Griffith et al., 2021; Láruson et al., 2022; St Clair et al., 2022; Varas-Myrik et al., 2022; Lind et al., 2024; reviewed in Capblancq et al., 2022). Such analyses could play a key role in restoration efforts by identifying populations or regions to be prioritized for *ex situ* collections, quantifying regions of connectivity to monitor for potential EAB movement, preserving genomic variation critical to adaptation under current and future environments, and designing pest resistance breeding programs.

*Fraxinus latifolia* is a dioecious tree species characteristic of riparian areas across much of the Pacific Northwest of North America. With a geographic distribution ranging from southern California, USA to British Columbia, Canada, it is one of the few deciduous tree species widespread across western coastal forests. It plays critical roles in regulation of light and water regimes in the region and is commonly used in restoration, particularly within wetland systems where it plays a primary successional role (Prive, 2016). While widespread, the impending threat and magnitude of mortality associated with EAB has garnered the species near-threatened conservation status globally according to the IUCN Red List of Threatened Species (Oldfield & Westwood, 2017). To date, no studies have been conducted describing the genomic structure of *F. latifolia* to ensure population genomic variation and adaptive evolutionary potential is considered within conservation and restoration management. Thus, *F. latifolia* provides an ideal case to evaluate factors influencing range-wide patterns of genomic variation to support proactive conservation applications to expanding *ex situ* collections and establishing resources foundational to an EAB-resistance breeding program.

Leveraging the power of landscape genomics for a widespread species at risk enables identification of regions and populations of greatest conservation concern and can improve selection of populations for expanding *ex situ* collections and breeding programs. Given the immediate risk of EAB to larger parts of the *F. latifolia* range, there is a need for proactive conservation. While extensive preparation and response efforts (Bliss-Ketchum et al., 2021) have been in place prior to EAB’s detection in Oregon, USA and British Columbia, Canada, this study represents the first application of genomic data to conservation and restoration for *F. latifolia*.

Using high-throughput sequencing across populations spanning the distribution of *F. latifolia,* we will (*i*) characterize contemporary genomic variation within and among populations of *F. latifolia*, (*ii*) evaluate gene flow and identify potential corridors of connectivity across populations, (*iii*) assess contemporary genotype-environment associations and predict capacity to adapt to change, and (*iv*) identify populations or regions of greatest conservation concern. These data and analyses will provide a valuable reference for long-term genomic monitoring across the species’ range that will be critical as EAB continues to expand its range, and baseline understanding of connectivity and adaptation needed to identify germplasm to target for current and future conservation, preservation, and restoration efforts.

## Materials and Methods

### Population sampling and DNA extraction

Between the autumn of 2021 and spring of 2022, leaf tissue was collected from individuals spanning nearly the entire natural distribution of *Fraxinus latifolia* (Oregon ash; Figure 1A; Table S1). 61 populations were sampled across the species’ distribution from California, Oregon, Washington, and British Columbia, for a total of 1,083 individuals. When possible, at least 20 individuals were sampled per population. Fresh leaf tissue was dried on silica gel and approximately 20 mg of dry tissue was used for DNA extraction. DNA was extracted locally using a Macherey-Nagel NucleoSpin Plant 2 extraction protocol or out-sourced via a high-throughput modified CTAB protocol by Ag-Biotech (Monterey, CA, USA). The concentration and purity of extracted DNA was quantified for each sample using a NanoDrop 1000 Spectrophotometer (Thermo Fisher Scientific; Waltham, MA, USA) to ensure all samples had a concentration of at least 2.0 ng/⎧L and purity ratios, per 260/280, of 1.21 to 2.09 (average 260/280 = 1.93).

**Figure 1.**
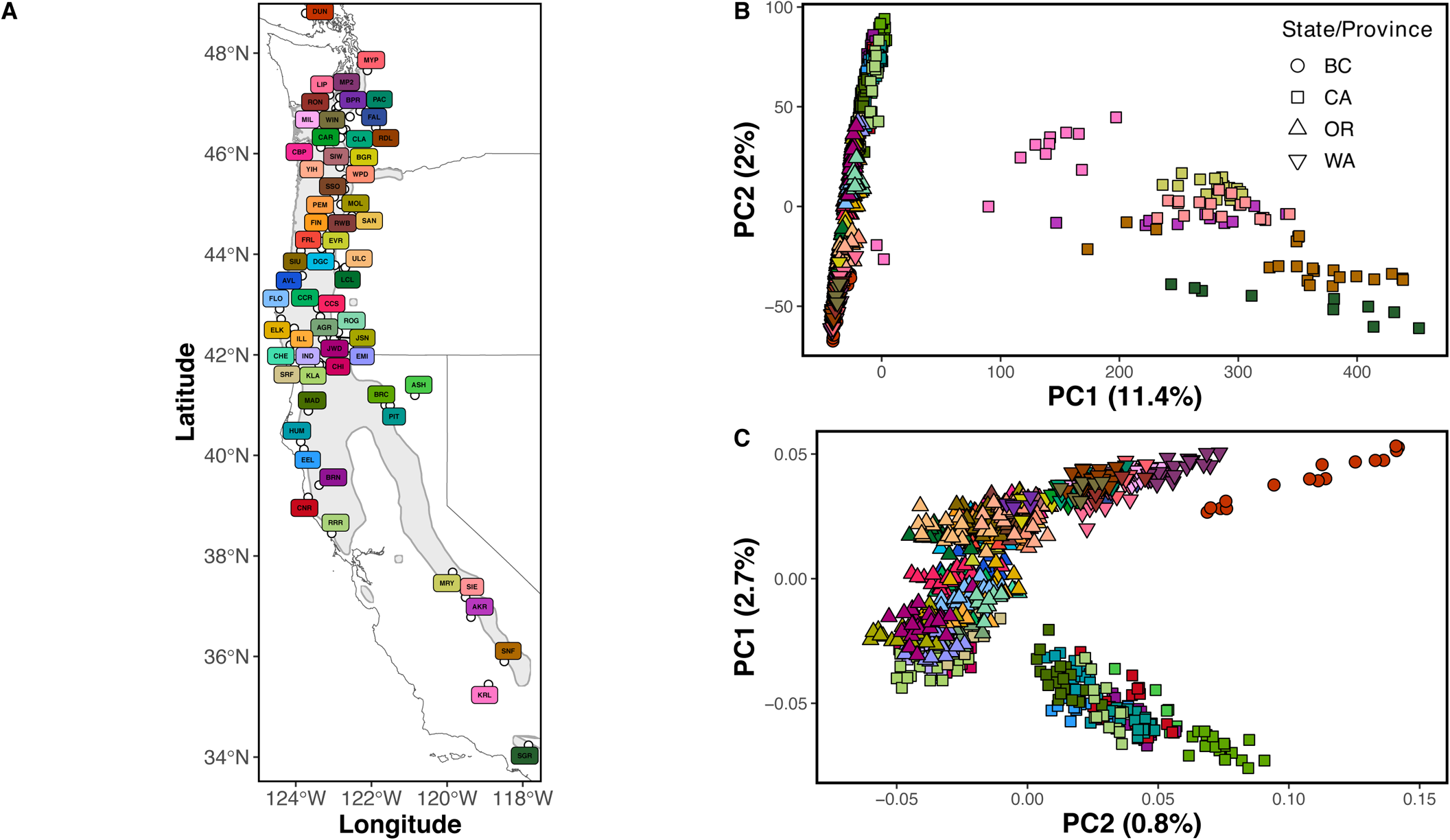
Map of sampled populations (A), PCA results for SNPs from all samples (B), and PCA results for all samples excluding southern populations (C). The southern samples were significant outliers in both the SNP PCA (B) and allelic proportions PCA (Fig. S1) and were thus excluded from a second SNP PCA (C) and all subsequent analyses. The x-axis of (C) is PC2, versus PC1, to highlight the geographic structure of the populations and match the orientation of the map, with the northernmost population being high on the y-axis, and the southernmost populations being low. Color of population markers in (A) is consistent across all figures and listed in Table S1. The gray outline and shaded region in (A) represent the range of *Fraxinus latifolia*, per Little, 1971 and acquired from databasin.org.

### Genomic library preparation and ddRADseq

Genomic libraries were prepared using a double-digest restriction-site associated DNA sequencing (ddRADseq) protocol per Parchman et al. (2012). Genomic DNA was digested using *Eco*RI and *Mse*I endonucleases (New England BioLabs, Inc). T4 DNA ligase (New England BioLabs, Inc) was then used to ligate uniquely barcoded Illumina adaptors (576 unique barcodes, each matching a sample) to the *Eco*RI cut sites of fragments, and standard Illumina adaptors to the *Mse*I cut sites. Barcoded fragments were amplified via PCR using Iproof DNA polymerase (BioRad; Hercules, CA USA). PCR-amplified genomic libraries were pooled and sent to the Genomic Sequencing and Analysis Facility (GSAF; Austin, TX, USA) for size selection of fragments within the range of 350 to 450 bp using a Pippin Prep quantitative electrophoresis unit (Sage Science; Beverly, MA, USA). Single-end sequencing of 100bp read lengths was performed on two lanes of an Illumina NovaSeq 6000 platform using S2 chemistry.

### De novo assembly and SNP calling

To detect and discard potential contaminants, such as *Escherichia coli*, *PhiX*, or Illumina-associated oligonucleotides we first used the ‘Tapioca’ pipeline (http://github.com/ncgr/tapioca). Demultiplexing of sequence files by individual was performed using a custom Perl script, which was also used to correct barcode sequencing errors, trim cut sites, and barcoded oligo-associated bases. Reads were filtered and assembled *de novo* using ‘cd-hit-est’ (Fu et al., 2012) following procedure detailed in ‘dDocent’ v2.7.8 (cutoffs: individual = 8, coverage = 6; clustering similarity: -c = 0.96; Puritz et al., 2014). This step generated a set of contig consensus sequences representing a reference of genomic regions sampled by the reduced representation sequencing data. Demultiplexed reads were mapped to the reference assembly using ‘bwa-mem’ v0.7.17 using default parameters (Li, 2013). Sequence variants were identified using ‘bcftools’ v1.9 (Danecek et al., 2021) and subsequent filtering was completed using ‘vcftools’ v0.1.16 (Danecek et al., 2011). Biallelic single nucleotide polymorphisms (SNPs) present within at least 90% of the sequenced reads across all samples were retained. These SNPs were further filtered to one per reference contig to reduce effects of linkage disequilibrium. Additional filtering included cut-offs of minor allele frequency ≥ 2%, mean read depth across samples ≥ 5 and < 15, and alternate allele call quality ≥ 999. SNPs with excessive coverage were removed and only those with two alleles present were retained to ameliorate potential genotyping bias from mis-assembly of paralogous genomic regions (Hapke and Thiele, 2016; McKinney et al., 2018). As mis-assembly of paralogous genomic regions can lead to an abnormal excess of heterozygosity, SNP-specific *F*_IS_ was calculated and only SNPs with *F*_IS_ < –0.5 were retained (Hohenlohe et al., 2013; McKinney et al., 2017). Individuals missing more than 40% of identified SNPs were removed from subsequent analyses (n = 30) resulting in a total of 998 individuals sequenced across 42,759 SNPs for inclusion in analyses below.

### Analysis of population genomic structure and diversity

To assess range-wide genomic structure of *F. latifolia,* a PCA was performed including all populations using the *snpgdsPCA* function within the ‘SNPRelate’ v1.34.1 R package (Zheng et al., 2012). In the southern portion of the *F. latifolia* distribution, hybridization with *F. velutina* Torr. has been reported and could involve production of tetraploids. (Taylor, 1945; Munz and Laudermilk, 1949; Twisselmann, 1967). Thus, to assess the potential for polyploidy, ploidy level variation was evaluated using the ‘gbs2ploidy’ v1.0 R package (Gompert & Mock, 2017) in R version 4.4.0 (R Core Team, 2024). See **Supplemental Methods** for more details. Samples from the southern terminus of the range (six populations, 92 individuals) were identified as outliers in principal component space, indicating substantial genomic differentiation in addition to evidence of polyploidy (see **Supplemental Results** for more details). Thus, all downstream analyses were performed excluding these individuals. To further characterize genomic structure and estimate admixture coefficients, a *structure* analysis was performed using ‘fastStructure’ (Raj et al., 2014) as implemented in the ‘structure-threader’ suite (Pina-Martins et al., 2017). The best-supported *k* -value was used to assess the number of genomic clusters (k = 2-6) using the ‘chooseK.py’ script from ‘fastStructure’ based on the remaining individuals 906 sampled across the species’ distribution.

Nucleotide diversity (π; (Nei & Li, 1979) and Watterson’s θ (Watterson, 1975) were calculated to describe population-specific variation in the distribution of polymorphisms. In addition, Tajima’s *D* (Tajima, 1989) was used to evaluate the mean number of pairwise differences across segregating sites for each population. This approach uses the distribution of rare alleles to indicate where and how different selective and demographic events may have shaped genomic variation. Analyses were implemented in ‘ANGSD’ v0.923 (Korneliussen et al., 2013, 2014) using methods that incorporate genotype uncertainty as this limits the potential impact of errors associated with mis-called genotypes where using low-coverage sequence data (Korneliussen et al., 2013). The folded site allele frequency likelihoods were estimated with *realsfs,* which calculated genotype likelihoods using the setting “GL 1” estimated from the model implemented in ‘samtools’ to obtain the likelihood of the folded site frequency spectrum (SFS). Nucleotide diversity was estimated using *doThetas 1* and *thetastat* commands using SFS likelihoods as priors, respectively, for each SNP across the reference assembly and averaged the measures for each population.

To estimate levels of inbreeding within populations, inbreeding coefficients (*F*_IS_) were calculated using the *gl.report.heterozygosity* function in the R package ‘dartR’ v2.9.7 (Mijangos et al., 2022) and relatedness within populations, quantified via identity-by-descent, was evaluated using the R package ‘SNPRelate’ v1.34.1’ was used (Zheng et al., 2012). Using the *snpgdsIBDMoM* and *snpgdsIBDMLE* functions, which use method of moments (MoM) (Purcell et al., 2007) and maximum likelihood estimation (MLE) (Milligan, 2003; Choi et al., 2009) methods, respectively, kinship coefficients for each possible pair within a population were compared. Finally, to estimate contemporary effective population size (N_e_), the linkage disequilibrium effective population size (LD-N_e_) was estimated for each population using the *ldNe* function from the ‘StrataG’ R package v2.5.01 (Archer et al., 2017) per methods described by Waples et al. (2016), with a minimum allele frequency of 0.05 and default parameters. Effective population size provides a valuable population parameter that predicts the rates of genetic drift and can relate to levels of inbreeding (Franklin et al., 1980; Braasch et al., 2021; reviewed in Waples, 2022, 2024; Gargiulo et al., 2024).

To quantify genomic differences among populations, SNP-specific pairwise estimates of Hudson’s *F*_ST_ (Hudson et al., 1992) and Nei’s *D* (Nei, 1972) were calculated from allele frequencies. Hudson’s *F*_ST_ estimation was selected as it is consistent across sampling schemes and independent of sample composition (Bhatia et al., 2013).

### *Estimates of connectivity for* F. latifolia

To quantify and visualize population connectivity across the range of *F. latifolia*, Estimated Effective Migration Surfaces (‘EEMS’; Petkova et al., 2016) was used to estimate species’ effective migration surfaces. The ‘Plink’ v2.0 (Chang et al., 2015) function “--make-bed” was used to convert the processed VCF to a BED file, allowing for non-human chromosome counts with the “--allow-extra-chr” flag. A difference matrix using the “mean allele frequency” imputation method was generated using the bed2diffs_v2 function from the ‘EEMS’ R scripts (https://github.com/dipetkov/eems; Petkova et al., 2016). Multiple ‘EEMS’ runs were performed to evaluate the goodness of fit of models derived using different “nDemes” parameters. Models were developed using “nDemes” starting at 100 and increasing by 100 each run until AICc scores no longer improved and observed demes did not increase. The best performing model was then selected using AICc scores. Models were generated using 25 million MCMC iterations, a burn-in of 10 million iterations, and 9,999 removed between samplings.

In addition, expected heterozygosity (*H*_E_) and observed heterozygosity (*H*_O_) were calculated using the *gl.report.heterozygosity* function in the R package ‘dartR’ v2.9.7 (Mijangos et al., 2022). *H_O_* was interpolated across the species’ range to identify regions across the landscape that may be targets for additional *ex situ* collections. Interpolations were performed using the inverse distance weighting (IDW) method as implemented in the R package ‘spatstat’ v3.0-3 (Baddeley et al., 2015).

### Quantifying the influence of geography and environment on genomic variation

The multivariate relationship between geography, environment, and genomic variation was quantified using 23 climatic variables averaged between 1991 and 2020 (Table S1) extracted from ClimateNA v7.41 (Wang et al., 2016) at a 1km^2^ resolution based on longitude and latitude of population origins (Tables S1). Elevation for each population was extracted using the ‘elevatr’ R package v0.4.5 (Hollister et al., 2023). Collinearity across climatic variables was assessed using the *cor* function in the base R package ‘stats’ to calculate Pearson correlations among climatic variables to limit potential redundancies. Variables with a correlation coefficient greater than |0.75| were excluded from further analyses. Seven climatic variables were retained following multicollinearity reduction, including frost-free period (FFP), mean annual precipitation (MAP), mean annual temperature (MAT), precipitation as snow (PAS), relative humidity (RH), and continentality (TD, temperature difference between mean warmest month temperature and mean coldest month temperature; Table S1).

Both isolation-by-distance (ie., IBD; population genomic distances associated with geographic distances) and isolation-by-environment (ie, IBE; accumulation of genomic differences between populations from distinct environments), were evaluated using partial mantel tests with the *mantel.partial* function from the R package ‘vegan’ v2.6-4 (Oksanen et al., 2022). The tests were performed using the Nei’s *D* genetic distance metric, Euclidean environmental distances per the *dist* function of base R, Haversine geographic distances converted to kilometers per the *distHaversine* function of the ‘geosphere’ v1.5-18 R package (Hijmans et al., 2022), 999 permutations, and Pearson’s product-moment correlation. A Mantel test was also used to test for correlation between geographic and environmental distances.

To test for genotype-environment associations and the impact of climate on population genomic structure, a redundancy analysis (RDA) was performed, with latitude and longitude included to represent population geographic structure (see **Results**). Prior to analysis, elevation and climate variables were scaled using the base R *scale* function and SNP data was converted to minor allele counts using ‘vcftools’ (--012 output format). The *rda* function from the R package ‘vegan’ v2.6-4 was used to perform RDA using uncorrelated environmental variables, including latitude and longitude to represent geography, and elevation, frost-free period (FFP; days), mean annual precipitation (MAP; mm), mean annual temperature (MAT; °C), relative humidity (RH; %), and continentality (TD; °C) to represent the environment. The ‘vegan’ v2.6-4 function *varpart* was used to perform a variance partitioning analysis based on the RDA model to evaluate geographic distance, environmental distance, and their individual and combined effect on genomic distance among populations.

### Predicting genomic offset under climate change

A generalized dissimilarity model (GDM) was generated to model changes in genomic distances along climatic gradients and forecast genomic changes required to maintain genotype-environment relationships under future climate conditions (i.e., genomic offsets; Fitzpatrick & Keller, 2015). Genomic distances were calculated using the *dist.genpop* function of the ‘adegenet’ R package (Jombárt, 2008) for all SNPs, and climatic variables previously used for RDA and IBE were used as the environmental matrix (FFP, MAP, MAT, PAS, RH, and TD). The *gdm* function within the R package ‘gdm’ v1.5.0-9.1 (Fitzpatrick et al., 2022) was used to fit the generalized dissimilarity model to the genomic distance and climatic data. Ensemble projections associated with future climate averages between the years 2041-2070 using three climate change emission scenarios: ssp245 (a less severe change in climate assuming a decrease in carbon output), ssp370 (a very likely scenario with continued carbon output), and ssp585 (a “worst case scenario” model assuming an increase in carbon output; Mahony et al., 2022) were used for GDM. Ensemble models were used to account for potential spatio-temporal variation associated with individual models to increase confidence in climate change projections (Mahony et al., 2022; reviewed in Hausfather et al., 2022). The *gdm.transform* function was used to rescale climatic variables based on variable importance in the GDM. Transformed climatic variables were then used as input for a PCA using the base R prcomp function with a random sampling of 50,000 raster cells. The first three principal components were mapped to RGB color values which were combined for each cell and projected onto the landscape to evaluate how future climate projections will contribute to genomic offset across the species range.

## Results

### Population genomic structure

After mapping and variant calling, we initially retained 1,280,333 biallelic SNPs across the 1,083 individuals. Following stringent filtering, 42,759 SNPs remained for 998 individuals (average sequencing depth = 7.92x) representing 61 populations across the range of *F. latifolia* (Figure 1A; Table S1 for population sampling information).

An initial principal components analysis using genomic data for all individuals revealed two distinct clusters largely differentiating the northern and southern distribution of sampled populations. PC1 explained 10.34% of the variance distinguishing two distinct clusters and PC2 1.96% of the variance (Fig. 1B; Fig. S1B). One cluster comprised the majority of individuals sampled from British Columbia, Washington, Oregon, and the northern coast of California, while the second cluster included individuals sampled from the southern range of the Sierra Nevada mountains and disjunct southwestern populations. Given the gap in sampling between the northern and extreme southern groups, uncertainty in the potential for hybridization between *F. latifolia* and *F. velutina* in the south, and evidence here (Fig. S2) and elsewhere (Taylor, 1945; Munz and Laudermilk, 1949; Twisselmann, 1967) for polyploidy in the southern part of the range, we limited subsequent landscape genomic analyses to 906 samples from 55 populations (∼85% of our samples), excluding these six genomically distinct populations (**Supplemental Methods, Supplemental Results**, Table S2). Subsequent analysis based on the *F. latifolia* populations ranging from British Columbia to northern California identified three clusters that largely followed a north to south latitudinal gradient indicated by a strong correlation between latitude and PC1 (r = 0.931; Fig. 1C).

*Structure* analysis identified *k* = 4 (marginal likelihood = -0.743; Fig. S3B) as the model that maximized marginal likelihoods of cluster assignment and *k* = 2 (marginal likelihood = - 0.744; Fig. S3B) as the number of clusters that best explain genomic structure within the data. For *k* = 2, a north-to-south gradient was evident, with the British Columbia, Canada population comprising a ‘northern’ cluster, and samples exhibiting mixed ancestry across clusters within northern Oregon along the Willamette River Valley differentiated from southern populations of the southern cluster in southern Oregon and California (Fig. S3A). The north-to-south gradient was prominent at *k*=4, with clusters representing (*i*) the British Columbia, Canada population, (*ii*) Washington to central Oregon, USA populations, (*iii*) southwest Oregon and northwest California, USA populations adjacent to Oregon, (*iv*) and all other California, USA populations (Fig. S3A).

### Population genomic diversity analyses

Genome-wide estimates of nucleotide diversity (π) were low across all populations. The northernmost population in British Columbia (DUN) exhibited the lowest nucleotide diversity (3.48x10^-3^±1.49x10^-5^). Populations at the northern limits and in smaller river valleys towards the eastern and western range extents exhibited reduced nucleotide diversity relative to those populations in the central river valleys of the species’ distribution. Estimates of π ranged from 3.48x10^-3^±1.49x10^-5^ (DUN, British Columbia, Canada) to 5.54x10^-3^±1.66x10^-5^ (PIT, California, USA), with a mean of 4.85x10^-3^ (Fig. 2A, Fig. S4A). Similar to π, Watterson’s θ values were low across all populations, exhibiting a similar latitudinal distribution, from 2.28x10^-5^±8.96x10^-6^ (DUN, British Columbia, Canada) to 5.13x10^-3^±1.23x10^-5^ (PIT, California, USA), with a mean of 4.20x10^-3^ (Fig. S4B). Tajima’s *D* values were negative for all but one population (CHE, Oregon, USA; Fig. S4C), indicating a potential recent population expansion after a genetic bottleneck. Tajima’s *D* ranged from -3.72x10^-2^±4.83x10^-5^ (HUM, California, USA) to 1.91x10^-3^±2.74x10^-5^ (CHE, Oregon, USA), with an average of -2.48x10^-2^.

**Figure 2.**
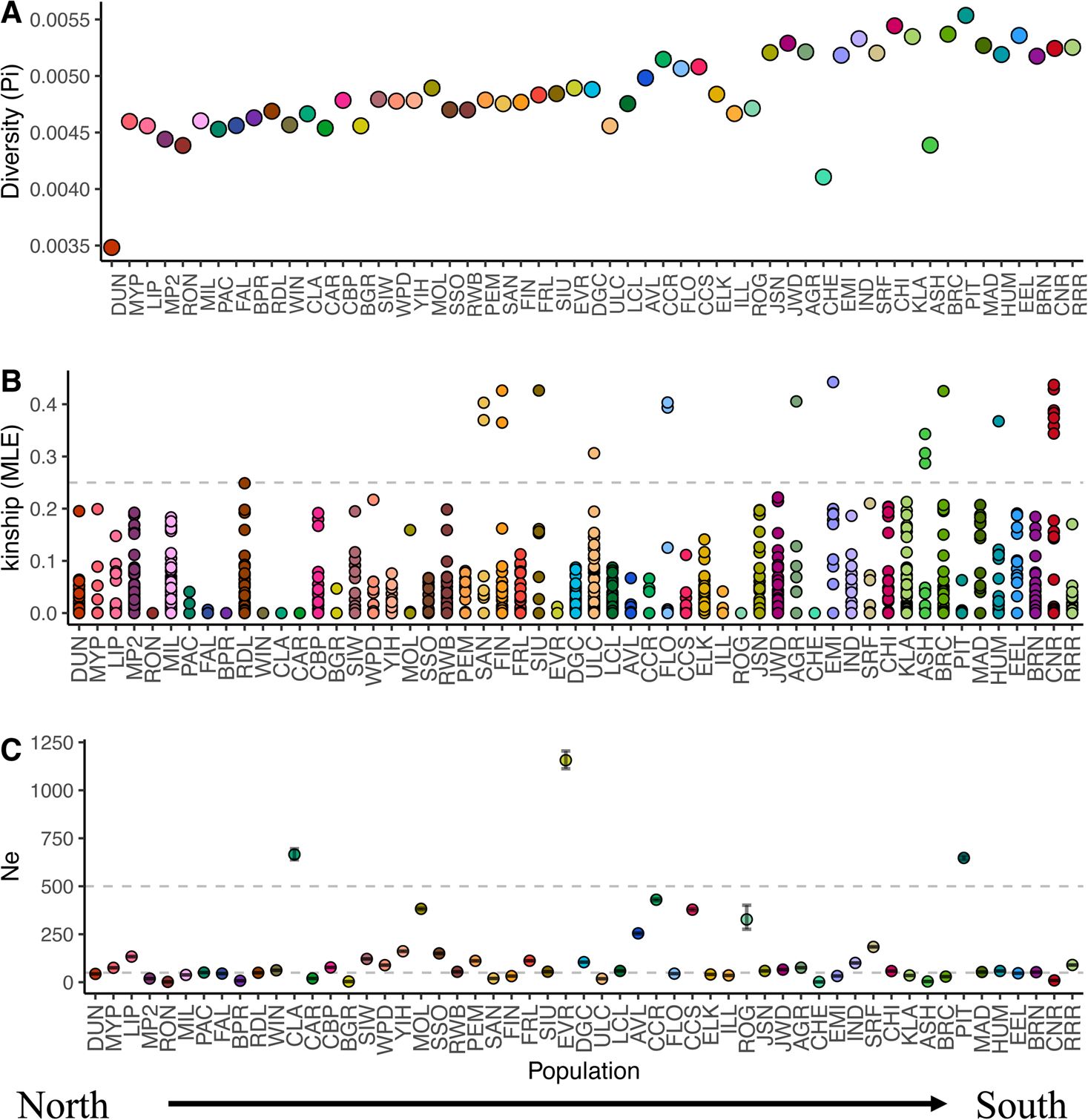
Dotplots visualizing nucleotide diversity (π; A), MLE relatedness estimates (B), and LD-N_e_ (C), per population, ordered by decreasing latitude. π estimates were generally very low across populations ranging from 3.48x10^-3^ and decreased with increases in latitude. Relatedness estimates were low, with means per population of MLE estimates ranging from 0 to 0.03, with similar results for MoM estimates. Horizontal lines in (B) designate the coefficient of full-sibs, 0.25. Points above these lines represent samples with kinship coefficients greater than 0.25 and are more closely related than full-sibs. Effective population size estimates were very low for all populations, with all but three falling below 500 (PIT = 650, CLA = 670, and EVR = 1,160). Horizontal lines are at 50 and 500, which are commonly accepted thresholds for N_e_ to combat inbreeding depression (50) and genetic drift (500).

Estimates of *F*_IS_ were generally low, ranging from -0.151 (DUN, British Columbia, Canada) to 0.056 (SRF, California, USA) with a mean of 0.014±0.035, suggesting outbreeding within populations typical of a dioecious system. Identity-by-descent analysis also produced very low relatedness estimates. Only 0.3% (24 of 7,810) of pairwise kinship coefficients were greater than 0.25. The majority of kinship coefficients (7,786 of 7,810 comparisons) were very low (kinship coefficient < 0.25). Mean MoM estimates of relatedness per population ranged from 0 (no identity-by-descent among compared samples; RON and WIN, Washington, USA) to 0.114 (DUN, British Columbia, Canada), with a mean estimate of 0.016. Mean MLE estimates of relatedness per population ranged from 0 at seven populations to 0.03 (ASH, California, USA), with a mean estimate of 6.09x10^-3^ (Fig. 2B). Estimates of LD-N_e_ were generally low, ranging from 1.45 (CHE, Oregon, USA; *n* = 4) to 1,160 (EVR, Oregon, USA; *n* = 19), with a mean estimate of 128 ± 201 across all populations (Fig. 2C; Table S3). Only three populations exceeded N_e_ = 500 (PIT = 650, CLA = 670, and EVR = 1,160) and greater than twenty populations had N_e_ < 50, indicating limited evolutionary potential species-wide.

Genomic differentiation among the populations was moderate with low *F*_ST_ and Nei’s *D* values. *F*_ST_ values ranged from 0.030 (CCR, Oregon vs. CCS, Oregon) to 0.242 (CHE, Oregon, USA vs. DUN, British Columbia, Canada), with a mean of 0.078 (Fig. S5) and Nei’s *D* values ranging from 9.57x10^-3^ (CCR, Oregon, USA vs. CCS, Oregon, USA) to 0.094 (CHE, Oregon, USA vs. DUN, British Columbia, Canada), with a mean of 2.83x10^-2^ (Fig. S5).

### *Estimates of connectivity across the range of* F. latifolia

‘EEMS’ was used to identify departures from isolation-by-distance (IBD) to detect potential barriers to gene flow or areas with greater rates of effective migration than expected given geographic distance and to visualize population connectivity over the range *of F. latifolia* (Fig. 3). A total of seven ‘EEMS’ runs were performed, with “nDemes” ranging from 100 to 700, as likelihood scores and observed demes per nDeme no longer improved at higher nDeme values. Per likelihood metrics, the fifth run, with a demes grid of 500 Demes, was the best-fit model. Population connectivity, per estimated migration rates (*m*), was highest among demes occurring along major river systems in Oregon and northern California, indicating greater potential gene flow and genomic similarity than expected given their geographic distance. Estimates of effective migration rates were much lower outside of major river systems, indicating much higher genomic distances than expected given their geographic distances, suggesting the larger, more open riparian habitats serve as important corridors of gene flow for *F. latifolia* populations.

**Figure 3.**
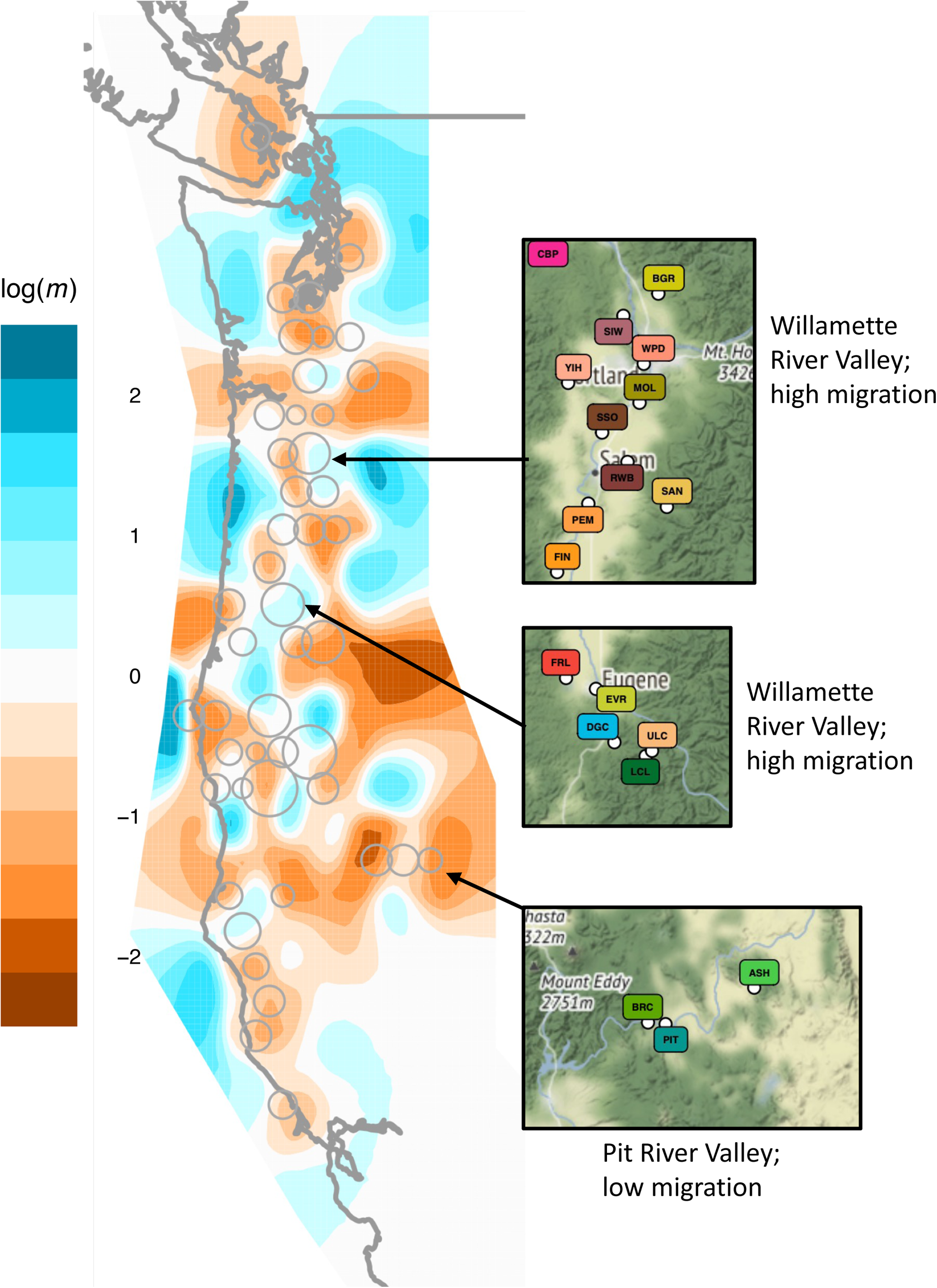
Estimated effective migration surface (‘EEMS’) visualization for *F. latifolia*. Differences in migration rates (*m*; on a log10 scale) and deme connectivity that were significantly higher than the overall average rates are represented in blue (*i.e.*, corridors of gene flow) and significantly lower than the overall average rate (*i.e.*, barriers to gene flow) are represented in brown. Demes used in the estimation of the migration surface are represented by dark gray circles, with the size of the circle corresponding to the number of samples within the deme. Overall, deme connectivity, and thus effective migration, was greatest along major river systems, such as Willamette River in Oregon. River systems outside of the central valleys of the species range featured less connectivity and effective migration (eg., the Pit River system).

Expected heterozygosity, *H*_E_, and observed heterozygosity, *H*_O_, were estimated to evaluate genomic diversity across corridors of connectivity. Estimates of *H*_E_ ranged from 0.182±0.278 (CHE, Oregon, USA) to 0.248±0.194 (KLA, California, USA) with a mean of 0.220±0.020. Estimates of *H*_O_ ranged from 0.143±0.189 (CHE, Oregon, USA) to 0.244±0.165 (CHI, California, USA) with a mean of 0.226±0.014. Greatest interpolated *H*_O_ (Fig. S6) was observed within populations occurring spanning large river systems central to the species’ distribution and coincides with predicted corridors of gene flow estimated from ‘EEMS’ (Fig. 3). Populations outside of the central valley systems, such as DUN, British Columbia, Canada and CHE, Oregon, USA, exhibited much lower *H*_O._ *H*_E_ and *H*_O_ were similar on average (mean *H*_O_/*H*_E_ = 0.974±0.048), although regionally several populations had much lower observed than expected heterozygosity (*H*_O_/*H*_E_ < 0.9; CHE = 0.783, DUN = 0.837, ASH = 0.872, and BPR = 0.892) indicating that these populations may experience decreased gene flow, and could be more susceptible to drift.

### Quantifying the relationship between environment, geography, and genomic variance

Variance partitioning indicated that the combined effects of geography and environment explained 29.3% of the genomic variance among populations (residuals = 0.685; Fig. 4D). Independently, geography and environment respectively accounted for 0.9% and 1.2% of genomic variance. Both IBD and IBE had statistically significant influences on genomic variation, however, their joint influence was greatest. Partial Mantel tests were statistically significant for IBD with IBE removed (*r* = 0.337, p-value = 0.001; Fig. 4A) and IBE with IBD removed (*r* = 0.208, p-value = 0.027; Fig. 4B), indicating that while both geography and environment influence population genomic structure of *F. latifolia*, geography alone explains more than environment. The Mantel test for environmental distance by geographic distance was also statistically significant (r = 0.182, p-value = 0.004; Fig 4C).

**Figure 4.**
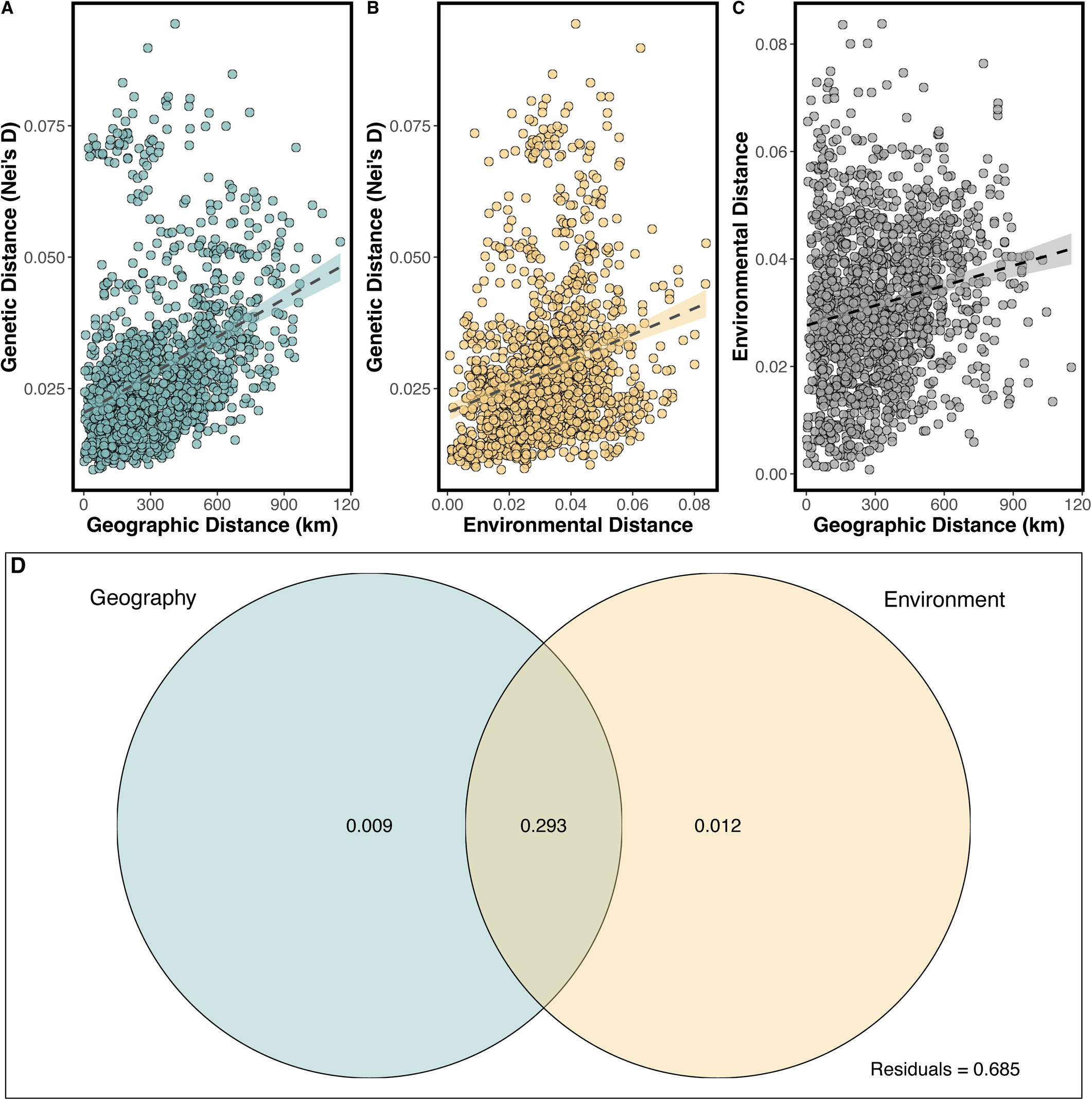
Results of IBD, IBE, and IBD-IBE analyses. Positive relationships were identified with Nei’s D for both geographic distance (A; r = 0.337, p-value = 0.001), environmental distance (B; r = 0.208, p-value = 0.027), environmental distance by geographic distance (C; r = 0.182, p-value = 0.004), and Venn diagram from *varparts* analysis showing the variance explained by geography and environment (D).

Environmental and distance variables that contribute to genomic structure were identified using RDA. RDA1 explained 51.75% of genomic variance (Fig. 5), with latitude, mean annual temperature (MAT), and relative humidity (RH) having the highest predictive loadings (eigenvectors). RDA2 explained 10.97% of variance with frost-free period (FFP) and continentality (TD; Fig. 5B) having the greatest predictive loadings. RDA3 explained 7.87% of variance with longitude (Fig. 5C) having the greatest predictive loading. Among northern populations (British Columbia and Washington) genomic structure was best explained by MAT (RDA1) and FFP (RDA2), while the regional genomic cluster including Oregon, and northern California populations was best explained by reduced MAT and greater RH (RDA1) and reduced continentality (RDA2). Southern California populations exhibited higher MAT on RDA1 and greater FFP (RDA2) relative to other regions. Overall, climatic gradients emphasize regional differences in genetic variance across the species’ range that are likely partially shaped by selection.

**Figure 5.**
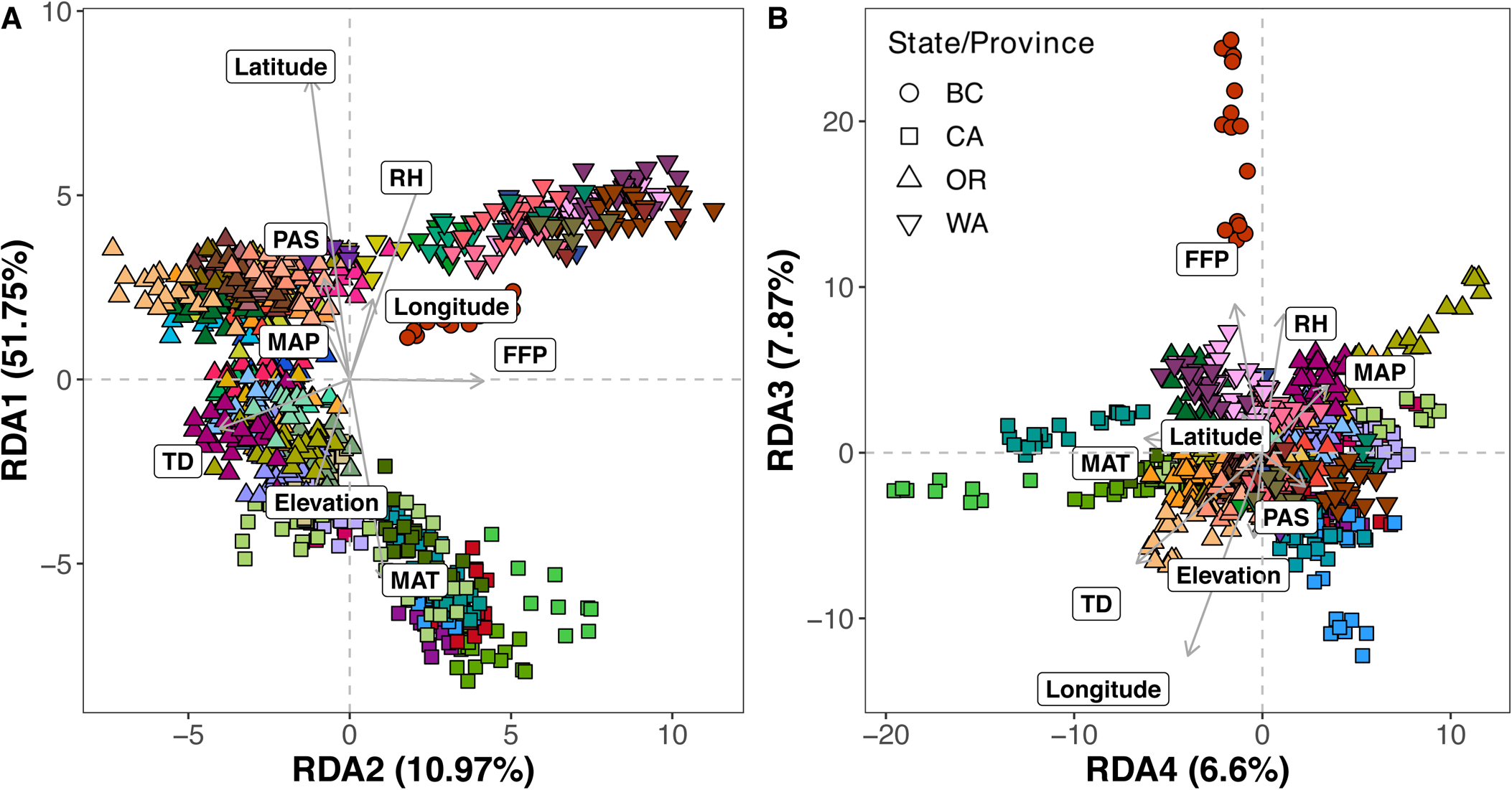
Population genomic structure was strongly associated with climatic variation across the range. RDA1, primarily comprising latitude and mean annual temperature (MAT), accounted for 51.75% of variances explained by the RDA model. RDA2, comprising largely frost-free period (FFP) and continentality (TD), explained 10.97% of variance.

### Predicting genomic offset under climate change

A generalized dissimilarity model was used to model changes in genomic distances along climatic gradients and to predict “home vs. away” genomic offsets, based on the uncorrelated climatic variables used in the RDA and variance partitioning analyses (FFP, MAP, MAT, PAS, RH, and TD). The final GDM included five informative climatic variables: FFP, MAT, MAP, RH, and TD. PAS was uninformative in GDM development. Evaluating different climate projection models indicated that ssp245, an optimistic scenario, exhibited predicted offset values from 0.087 to 0.130 (mean = 0.105), while ssp370, a very likely scenario exhibited values from 0.087 to 0.131 (mean = 0.106), and ssp585, a ‘worst case scenario’ exhibited values from 0.087 to 0.131 (mean = 0.108). Based on PCA, over much of the range of *F. latifolia,* predicted changes to mean annual temperature contributed to greatest genomic offset projections (Fig. 6; Fig. S7). However, in higher elevation regions, relative humidity had a stronger influence on genomic offset projections (Fig. 6). All remaining climatic variables largely loaded on PC3, indicating greatest potential projected offset within the eastern portion of the species’ range where fewer populations currently occur. The lowest projected offset was predicted in the southern portion of the species’ range, particularly along the southwestern edges of the range, suggesting these populations may be “pre-adapted” to a warming climate and will experience reduced offset under global change. In contrast, northern and eastern regions had predicted genomic offsets of 0.1 or greater, indicating these populations may not have the genomic variation needed to respond to changing selective pressures.

**Figure 6.**
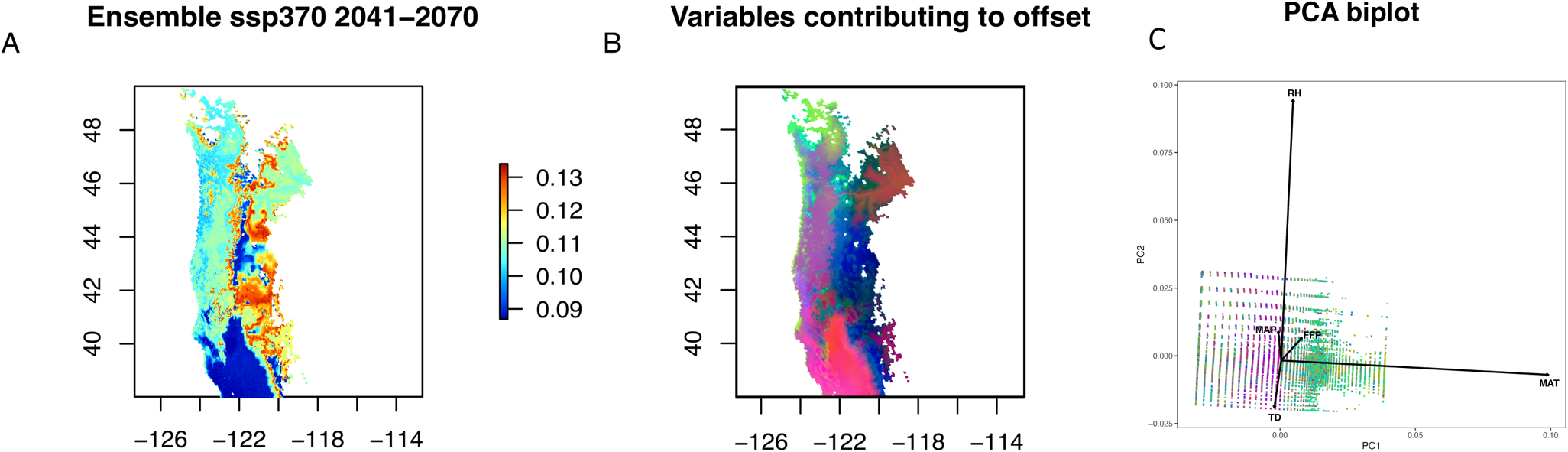
Visualization of predicted genomic offset (A visualizes predicted genomic offset under SSP370; other SSPs are visualized in Fig. S7), principal components of climatic variables most greatly contributing to predicted genomic offset (B), and bi-plot of principal components (C). Predicted genomic offsets were consistent across all shared socio-economic pathways (SSP) and ranged from 0.087 to 0.131 (mean = 0.106) for the most likely climate change scenario (ssp370; A). (B) Principal component 1 (red), primarily comprising MAT affects most of the range, principal component 2 (green), comprising largely RH, affect primarily the higher elevation areas along the primary valley systems within the species range, and principal component 3 (blue), comprising all other climatic variables, largely affects the eastern part of the range.

## Discussion

*Fraxinus latifolia* is foundational to wetland and riparian habitats of western North America. With the imminent threat of the emerald ash borer (EAB), understanding species-wide standing genomic variation has become critical to designing effective conservation and pest-resistant breeding programs. The majority of standing genomic variation for *F. latifolia* is structured along a north to south climate and latitudinal gradient, with an “abundant center” reflecting regions of nucleotide diversity and connectivity within central river valley systems and reductions in diversity among northern peripheral and disjunct populations in British Columbia, Oregon, and California, respectively. Despite evidence of connectivity, however, effective population size is generally low across all populations, which may indicate dioecy and patchy distributions have impacted evolutionary potential. Given predictions associated with genomic offsets, this may indicate that genomic changes required to maintain genotype-environment relationships under future forecasted conditions may be challenged and exacerbate risks associated with the invasion of EAB. However, these data provide a proactive path forward for defining conservation units needed to mitigate both the risk of climate change and invasion by EAB, ultimately laying the foundation for a genome-enabled conservation and EAB-resistance breeding program.

### Population genomic structure is driven by environment and geography

The genomic structure of *F. latifolia* largely follows a latitudinal gradient (Fig. 1C-2, Fig. S1C), with latitude a strong predictor of changes in allele frequencies. However, across the species’ distribution, genomic substructure indicates the combination of different neutral and non-neutral processes have likely influenced fine-scale genomic structure. Populations sampled in Washington and Oregon, which occurred primarily along central valley river systems of the Columbia River watershed, formed a large cluster comprising much of the genomic diversity within the species (Fig. 1, Fig. S3). Other areas, such as southern California, which included populations within the Pit River valley from which more geographically isolated populations were sampled, formed distinct clusters (Fig. S3). These results are consistent with an “abundant center” model reflecting increased species abundance towards the center of a distribution.

Indeed, regions of greatest connectivity (Fig. 3; Fig. S5) and observed heterozygosity (Fig. S6) were observed within the central portion of the species’ range. This pattern is likely driven by the riparian nature of the species (Franklin & Dyrness, 1973; Frenkel & Heinitz, 1987), as the populations along the largest river valleys experience greater connectivity. Similar latitudinal gradients in the distribution of genomic variation have been observed for other coastal trees of the Pacific Northwest. These patterns have been attributed to range expansion during the warming early Holocene (∼10,000 ybp) following the Pleistocene epoch (Pellatt et al., 2001, reviewed in Jaramillo-Correa et al., 2009), including *Picea sitchensis* (Bong.) Carr. (Mimura & Aitken, 2007; Mimura & Aitken, 2010), *Pinus albicaulis* Engelm. (Liu et al., 2016), *Pinus monticola* Dougl. ex. D. Don (Rehfeldt et al., 1984; Richardson et al., 2009; Kim et al., 2011), and *Populus trichocarpa* Torr. & A.Gray ex. Hook. (Geraldes et al., 2014). Furthermore, distributions of nucleotide diversity associated with latitude suggest that post-glacial expansion of *F. latifolia* likely followed from a single glacial refugia at the southern extent of the species’ distribution, supporting the “leading edge hypothesis” for post-glacial expansion (Soltis et al., 1997).

Both geography and environment strongly influenced the distribution of genomic variation across the species’ range. However, while both independently influence population genomic structure (Fig. 4), their combined influence had the greatest effect (29.3% variance explained; Fig. 4). This indicates that while both geographic and environmental distances do contribute significantly to population genomic structure, it is their combined influence that has the greatest predictive power in estimating genomic change.

### Central river valleys serve as corridors of gene flow

Population connectivity was greatest along major river systems within the center of the species distribution and lower along peripheral river systems. *F*_ST_ was generally low (mean *F*_ST_ = 0.078 ± 0.030; Fig. S5), indicating limited genomic differentiation among populations, with values in the range of those reported for other *Fraxinus* species (eg., for *F. excelsior* L., Italian populations *F*_ST_ = 0.049 (Ferrazzini et al., 2016), British populations *F*_ST_ = 0.025 for (Sutherland et al., 2010), and French populations *F*_ST_ = 0.016 (Sutherland et al., 2010)). However, the British Columbia, Canada (DUN) population (mean *F*_ST_ = 0.150 ± 0.024) and Chetco River, Oregon, USA (CHE) population (mean *F*_ST_ = 0.142 ± 0.025; Fig. S5) were genomically differentiated from all other populations. The DUN population represents the northern limit of the species’ range and is geographically isolated on Vancouver Island and disjunct from the mainland distribution (Fig. 1A). ‘EEMS’ indicated that the Salish Sea likely acts as an effective barrier to migration and gene flow between the mainland and island populations (Fig. 3). Similar patterns have been observed for populations of native tree species with island-mainland distributions (*Populus trichocarpa* (Gerlades et al. 2014) and *Pinus torreyana* Parry ex Carr. (Di Santo et al. 2022). The CHE population was sampled along the Chetco River in Oregon, USA. This river runs through rugged terrain that has likely acted as a physical barrier to gene flow, limiting potential influx of genomic variation from neighboring populations. While *F*_ST_ values were generally low, EEMS predicted regional heterogeneity in migration. Estimated effective migration rates appear highest along major river systems within the central valleys of the species range, such as Willamette river in Oregon, USA, and the Central Valley river systems of California (Fig. 3), while peripheral river systems exhibited lower rates (Pit River and Russian River valleys in California; Fig. 2). Regions with the greatest predicted effective migration reflect areas of greatest admixture also observed in the *structure* analysis (Fig. S3). Populations from southern Oregon and northern California, which occur primarily along large river systems within the Klamath-Siskiyou ecoregion that likely span a west-to-east region of connectivity (Eckert et al., 2008) formed a distinct cluster. Those populations in the north and south that were more geographically isolated from riparian habitats outside of central river valleys also formed relatively discrete clusters (Fig. S3).

The DUN population, geographically isolated on Vancouver Island, British Columbia, Canada was the most genomically distant of all populations (Fig. 1; Fig. S3-S4). This population comprised its own cluster within the *structure* analysis and was genomically differentiated from all populations (*F*_ST_ = 0.150 ± 0.024 and Nei’s *D* = 0.058 ± 0.087; Fig. S5), with extremely low nucleotide diversity (3.48x10^-3^±1.49x10^-5^; Fig. 2A). These results are likely attributed to past demographic changes, such as reduced connectivity, potential bottlenecks, and subsequent genetic drift following founding. These patterns are consistent with founding during the hypothesized northern expansion post-glaciation, followed by a long period of isolation (Soltis et al., 1997). Fossilized *Fraxinus* pollen grains found in the Saanich Inlet of British Columbia, Canada date back 9-10 kybp (Pellatt et al., 2001), suggesting this population likely was founded during this period, but has since become isolated and increasingly susceptible to loss of genomic variation through drift.

### Small effective population sizes in the presence of gene flow

Estimates of effective population size (LD-N_e_) were low across the range of *F. latifolia* (Fig. 2C). Most populations exhibited values well below N_e_ < 500, with over twenty populations with N_e_ < 50, suggesting the evolutionary potential of *F. latifolia* populations has likely been negatively influenced by fine-scale population substructure, drift, and a dioecious mating system. *Fraxinus latifolia* is a wind-pollinated dioecious plant species, and previous studies have suggested that habitat fragmentation across male and female plants can impact genetic exchange, influencing population genomic structure and reducing estimates of N_e_ (Bacles et al., 2005; Eisen et al., 2023). Furthermore, the combined influence of landscape heterogeneity and genomic structure associated with latitudinal gradients suggest that fine-scale population genomic substructure may have had a disproportionately negative influence on effective population size (Neel et al., 2013). Age stratification within populations has also been shown to negatively impact and bias effective population size estimates (Waples & Yokota, 2007; Waples, 2010; Waples et al., 2014). *Fraxinus latifolia* is a long-lived tree that generally does not reach reproductive maturity until approximately 30 years of age (Niemiec et al., 1995). However, while there is likely some degree of age stratification within our sampled populations, the majority of individuals were of reproductive age (Fig. S8), reducing potential biases associated with age stratification. Finally, biased sex ratios could lead to increased levels of inbreeding within populations, reducing effective population size (Ellstrand & Elam 1993; Frankham, 1995). Unfortunately, sex discrimination was not possible for all individuals, and therefore application of sex ratio estimates for associated with N_e_ estimation methods could not be used to make inferences of effective population size. However, estimates of relatedness were extremely low within populations (Fig. 2B), suggesting that N_e_ estimates were likely not biased by relatedness among individuals.

### Contemporary and future associations between genomic and climatic variation

In addition to strong associations with latitude, climatic gradients strongly influenced range-wide population genomic structure for *F. latifolia* (Fig. 5B; Fig. 6). Significant isolation by environment suggested increased population genomic differentiation with increasing environmental distance (r = 0.208, p-value = 0.027; Fig. 5B). Of climate variables, mean annual temperature (MAT) had the greatest predictive power on average of genomic differences across the species’ range (Fig. 5). Regional genomic differences among more northern populations of British Columbia, Canada and Washington, United States, were best predicted by temperature-related variables, including frost-free period (FFP) and continentality (TD; Fig. 5), distinguishing the cold and humid climates of northern British Columbia from the warmer, drier climates of California.

Genotype-environmental relationships currently maintained across the range of *F. latifolia* may be impacted by projections associated with future climatic conditions. Mean annual temperature (MAT) is predicted to change under all future climate change scenarios (IPCC, 2022). Based on the RDA, MAT was significantly associated with population genomic structure for *F. latifolia* (Fig. 5; Table S3-4). Evaluating genomic offset estimates, MAT was a primary predictor of future genomic offset across a large portion of the species’ range (Fig. 6; Fig. S7).

This suggests that as average temperatures rise across the range, populations will become more genomically distinct for genomic variation associated with adaptation to temperature gradients. Thus, as local climates warm, changes in MAT will impact predicted genotype-environmental associations. Furthermore, regional variation across the species’ range suggests populations are differentially sensitive to projected changes in temperature (eg., southern CA, USA populations are predicted to be less affected by increasing MAT than those in WA, USA). Similar patterns of increasing predicted genomic offsets at higher latitudes are seen in other western North American tree species, such as *Populus balsamifera* L. (Fitzpatrick & Keller, 2015) and *Pseudotsuga menziesii* var. *glauca* (Mayr) Franco (Lind et al., 2024). Together, these data suggest that potential mismatches between standing genomic variation and the genomic variation needed to adapt to changing environmental conditions may have disproportionate effects across the species’ range. However, these results should be interpreted with caution, as recent studies have shown that predicted genomic offsets may not necessarily represent “genomic vulnerability” and may require experimental results from common gardens to confirm predicted effects of changing environments on a given genotype (Lotterhos, 2024).

### Implications for conservation

The invasive EAB has decimated native *Fraxinus* populations across eastern North America and has now been observed in Forest Grove, Oregon, USA in 2022 with indications that it may be spreading locally and recently confirmed in British Columbia, Canada (April 2024). Where freezing temperatures have traditionally limited the spread of EAB, expected increases in temperature associated with climate change and increased frost-free periods suggest that resistance to invasion across the range of *F. latifolia* is likely reduced (DeSantis et al., 2013). Indeed, the increased risk of genomic offset posed by climate change, which may have fitness consequences associated with abiotic stress (Fig. 6; Fig. S7), puts *F. latifolia* at greater risk of local extirpations across its range. While estimates of population connectivity in the ‘abundant center’ along large river valleys central to the species’ range, provide potential corridors of movement for pollen and seed flow, it is clear that these regions may also facilitate the spread of EAB and should be monitored.

Since 2019, efforts have been in place to conserve germplasm *ex situ* with a long-term goal to assess resistance to EAB in *F. latifolia* populations leveraging knowledge from breeding programs established in the eastern United States (Sniezko, 2020; Bliss-Ketchum et al., 2021). Through 2023, seeds from over 350 maternal parent trees have been collected and preserved at the USDA Forest Service’s Dorena Genetic Resource Center (Cottage Grove, OR, USA) and the National Seed Laboratory (Dry Branch, GA, USA) providing a baseline of material to establish progeny trials and breed for resistance. While these collections will aid in establishing an EAB-resistance breeding program, the collections do not cover genomic variation spanning the range of *F. latifolia*. Given the results presented here, existing collections may lack genomic variation that represents neutral or adaptive genomic variation critical to long-term species evolution.

Thus, these data are invaluable to identifying regions to target additional *ex situ* collections for maintenance of germplasm needed for both progeny trials and long-term conservation. Climate change will likely exacerbate the risks to *F. latifolia,* as the combination of abiotic and biotic threats have already led to high mortality in several *Fraxinus* species in eastern North America, Russia, and Ukraine (reviewed in Sun et al., 2024). Conservation of genomic diversity will be key to the long-term conservation and restoration of *F. latifolia* and application of genomic data to monitoring provides a proactive means to mitigate potential threats associated with an invasive species. Ultimately, combining conservation and genomics approaches will facilitate breeding and screening for resistance, leveraging a genomics-driven pipeline for forest-pest management critical to long-term conservation, breeding, and restoration.

## Acknowledgements

This work was supported by a USDA National Institute of Food and Agriculture, McIntire Stennis project 1027676 to JAH. The authors would like to thank individuals from the Schatz Center for Tree Molecular Genetics, Department of Ecosystem Science and Management at Pennsylvania State University (M. Mailander and J. Beeman), the British Columbia Ministry of Forests (A. Yanchuk), the California Department of Forestry and Fire Protection (C. Lee), the US Dept. of Agriculture (J. Bronson, S. Johnson, S. Kolpak, and J. Watson), the USDA Forest Service (D. Cluck, K. Farris, P. Folsom, C. Harrington, T. Litmann, G. de Nevers, H. Prendeville, C. Starling, B. Willhite, and C. Winkler), Washington State University (G. Chastagner, S. Harvey, P. Shults, and K. Zobrist), the University of Washington (D. Peterson), the Washington state Center for Natural Lands Management (S. Freed), M. Habehicht, E. Man, and D. Pigott for providing samples that were used in this research. Any use of trade, firm, or product names is for descriptive purposes only and does not imply endorsement by the U.S. Government.

## Data Accessibility and Benefit-Sharing

Demultiplexed sequence data are to be uploaded to the National Centre for Biotechnology Information database and made publicly available upon acceptance of the manuscript. Scripts used to perform dDocent and ANGSD analyses are available at https://github.com/trevorfaske/FRLA_analyses. Scripts to perform downstream analyses are available at https://github.com/aemelton/OregonAshLandscapeGenomics.

## Author contributions

RS, TT, WW, and JAH contributed to planning and performing sample collections. AEM, TMF, TP, and JAH designed the research. TP performed ddRADseq genomic library preparation and sequencing. AEM and TMF performed the analyses. AEM wrote the initial draft of the manuscript. All authors contributed to the revision and finalization of the manuscript.

## References

Archer, F. I., Adams, P. E., & Schneiders, B. B. (2017). StrataG: An r package for manipulating, summarizing and analysing population genetic data. Molecular Ecology Resources, 17(1), 5–11. 10.1111/1755-0998.12559

Bacles, C. F. E., Burczyk, J., Lowe, A. J., & Ennos, R. A. (2005). Historical and contemporary mating patterns in remnant populations of the forest tree Fraxinus excelsior L. Evolution, 59(5), 979–990. 10.1111/j.0014-3820.2005.tb01037.x

Baddeley, A., Rubak, E., & Turner, R. (2015). Spatial Point Patterns: Methodology and Applications with R. London: Chapman and Hall/CRC Press.

Barstow, M., Oldfield, S., Westwood, M., Jerome, D., Beech, E., & Rivers, M. (2018). The Redlist of Fraxinus. BCGI. Richmond, UK.

Bhatia, G., Patterson, N., Sankararaman, S., & Price, A. L. (2013). Estimating and interpreting FST: The impact of rare variants. Genome Research, 23(9), 1514–1521. 10.1101/gr.154831.113

Bliss-Ketchum, L., Draheim, R., Hepner, M., & Guethling, O. (2021). Readiness and response plan for Oregon.

Bolte, C. E., Phannareth, T., Zavala-Paez, M., Sutara, B. N., Can, M. F., Fitzpatrick, M. C., Holliday, J. A., Keller, S. R., & Hamilton, J. A. (2024). Genomic insights into hybrid zone formation: The role of climate, landscape, and demography in the emergence of a novel hybrid lineage. Molecular Ecology, May, 1–16. 10.1111/mec.17430

Borrell, J. S., Zohren, J., Nichols, R. A., & Buggs, R. J. A. (2020). Genomic assessment of local adaptation in dwarf birch to inform assisted gene flow. Evolutionary Applications, 13(1), 161–175. 10.1111/eva.12883

Braasch, J. E., Di Santo, L. N., Tarble, Z. J., Prasifka, J. R., & Hamilton, J. A. (2021). Testing for evolutionary change in restoration: A genomic comparison between ex situ, native, and commercial seed sources of *Helianthus maximiliani*. Evolutionary Applications, 14(9), 2206–2220. 10.1111/eva.13275

Capblancq, T., Fitzpatrick, M. C., Bay, R. A., Exposito-Alonso, M., & Keller, S. R. (2020). Genomic Prediction of (Mal)Adaptation across Current and Future Climatic Landscapes. Annual Review of Ecology, Evolution, and Systematics, 51, 245–269. 10.1146/annurev-ecolsys-020720-042553

Carlson, S. M., Cunningham, C. J., & Westley, P. A. H. (2014). Evolutionary rescue in a changing world. Trends in Ecology and Evolution, 29(9), 521–530. 10.1016/j.tree.2014.06.005

Cavender, N., Westwood, M., Bechtoldt, C., Donnelly, G., Oldfield, S., Gardner, M., Rae, D., & McNamara, W. (2015). Strengthening the conservation value of *ex situ* tree collections. Oryx, 49(3), 416–424. 10.1017/S0030605314000866

Choi, Y., Wijsman, E. M., & Weir, B. S. (2009). Case-control association testing in the presence of unknown relationships. Genetic Epidemiology, 33(8), 668–678. 10.1002/gepi.20418

Conservation, S. H. (n.d.). Willamette Valley Conservation Study.

Danecek, P., Auton, A., Abecasis, G., Albers, C. A., Banks, E., DePristo, M. A., Handsaker, R. E., Lunter, G., Marth, G. T., Sherry, S. T., McVean, G., & Durbin, R. (2011). The variant call format and VCFtools. Bioinformatics, 27(15), 2156–2158. 10.1093/bioinformatics/btr330

Danecek, P., Bonfield, J. K., Liddle, J., Marshall, J., Ohan, V., Pollard, M. O., Whitwham, A., Keane, T., McCarthy, S. A., & Davies, R. M. (2021). Twelve years of SAMtools and BCFtools. GigaScience, 10(2), 1–4. 10.1093/gigascience/giab008

DeSantis, R. D., Moser, W. K., Gormanson, D. D., Bartlett, M. G., & Vermunt, B. (2013). Effects of climate on emerald ash borer mortality and the potential for ash survival in North America. Agricultural and Forest Meteorology, 178–179, 120–128. 10.1016/j.agrformet.2013.04.015

Di Santo, L. N., Hoban, S., Parchman, T. L., Wright, J. W., & Hamilton, J. A. (2022). Reduced representation sequencing to understand the evolutionary history of Torrey pine (*Pinus torreyana* Parry) with implications for rare species conservation. Molecular Ecology, 31(18), 4622–4639. 10.1111/mec.16615

Des Roches, S., Pendleton, L. H., Shapiro, B., & Palkovacs, E. P. (2021). Conserving intraspecific variation for nature’s contributions to people. Nature Ecology and Evolution, 5(5), 574–582. 10.1038/s41559-021-01403-5

Diaz-Martin, Z., Fant, J., Havens, K., Cinea, W., Tucker Lima, J. M., & Griffith, M. P. (2023). Current management practices do not adequately safeguard endangered plant species in conservation collections. Biological Conservation, 280(September 2022), 109955. 10.1016/j.biocon.2023.109955

Eckert, A. J., Tearse, B. R., & Hall, B. D. (2008). A phylogeographical analysis of the range disjunction for foxtail pine (*Pinus balfouriana*, Pinaceae): The role of Pleistocene glaciation. Molecular Ecology, 17(8), 1983–1997. 10.1111/j.1365-294X.2008.03722.x

Eisen, A. K., Semizer-Cuming, D., Jochner-Oette, S., & Fussi, B. (2023). Pollination success of *Fraxinus excelsior* L. in the context of ash dieback. Annals of Forest Science, 80(1). 10.1186/s13595-023-01189-5

Ellstrand, N. C., & Elam, D. R. (1993). Population genetic consequences of small population size: Implications for plant conservation. Annual Review of Ecology and Systematics, 24, 217–242. 10.1146/annurev.es.24.110193.001245

Félix-Valdez, L. I., Vargas-Ponce, O., Cabrera-Toledo, D., Casas, A., Cibrian-Jaramillo, A., & de la Cruz-Larios, L. (2016). Effects of traditional management for mescal production on the diversity and genetic structure of *Agave potatorum* (Asparagaceae) in central Mexico. Genetic Resources and Crop Evolution, 63(7), 1255–1271. 10.1007/s10722-015-0315-6

Ferrazzini, D., Monteleone, I., & Belletti, P. (2007). Genetic variability and divergence among Italian populations of common ash (*Fraxinus excelsior* L.). Annals of Forest Science, 64(2), 159–168. 10.1051/forest:2006100

Fitzpatrick, M. C., & Keller, S. R. (2015). Ecological genomics meets community-level modelling of biodiversity: Mapping the genomic landscape of current and future environmental adaptation. Ecology Letters, 18(1), 1–16. 10.1111/ele.12376

Fitzpatrick, M., Mokany, K., Manion, G., Nieto-Lugilde, D., & Ferrier, S. (2022). gdm: Generalized Dissimilarity Modeling. R package version 1.5.0–9.1. https://CRAN.R-project.org/package=gdm

Frankham, R. (1995). Effective population size/adult population size ratios in wildlife: A review. Genetics Research, 66(2), 95–107. 10.1017/S0016672308009695

Franklin, J. F., & Dyrness, C. T. (1973). Natural Vegetation of Oregon and Washington.

Franklin, I. R., Soulé, M. E., & Wilcox, B. A. (1980). Conservation biology: an evolutionary-ecological perspective. Sunderland, MA: *Sinauer Associates*.

Frenkel, R. E., & Heinitz, E. F. (1987). Composition and structure of Oregon ash (*Fraxinus latifolia*) forest in William L. Finley National Wildlife Refuge, Oregon. Northwest Science, 61(4), 203–212.

Fu, L., Niu, B., Zhu, Z., Wu, S., & Li, W. (2012). CD-HIT: Accelerated for clustering the next-generation sequencing data. Bioinformatics, 28(23), 3150–3152. 10.1093/bioinformatics/bts565

Gandhi, K. J. K., & Herms, D. A. (2010). Direct and indirect effects of alien insect herbivores on ecological processes and interactions in forests of eastern North America. Biological Invasions, 12(2), 389–405. 10.1007/s10530-009-9627-9

Gargiulo, R., Decroocq, V., González Martínez, S. C., Paz Vinas, I., Aury, J., Lesur Kupin, I., Plomion, C., Schmitt, S., Scotti, I., & Heuertz, M. (2024). Estimation of contemporary effective population size in plant populations: Limitations of genomic datasets. Evolutionary Applications, 17(5), 2023–2027. 10.1111/eva.13691

Geraldes, A., Farzaneh, N., Grassa, C. J., Mckown, A. D., Guy, R. D., Mansfield, S. D., Douglas, C. J., & Cronk, Q. C. B. (2014). Landscape genomics of *Populus trichocarpa*: The role of hybridization, limited gene flow, and natural selection in shaping patterns of population structure. Evolution, 68(11), 3260–3280. 10.1111/evo.12497

Gompert, Z., & Mock, K. E. (2017). Detection of individual ploidy levels with genotyping-by-sequencing (GBS) analysis. Molecular Ecology Resources, 17(6), 1156–1167. 10.1111/1755-0998.12657

Griffith, M. P., Cartwright, F., Dosmann, M., Fant, J., Freid, E., Havens, K., Jestrow, B., Kramer, A. T., Magellan, T. M., Meerow, A. W., Meyer, A., Sanchez, V., Santiago-Valentín, E., Spence, E., Sustasche-Sustache, J. A., Francisco-Ortega, J., & Hoban, S. (2021). *Ex situ* conservation of large and small plant populations illustrates limitations of common conservation metrics. International Journal of Plant Sciences, 182(4), 263–276. 10.1086/713446

Hamilton, J., Flint, S., Lindstrom, J., Volk, K., Shaw, R., & Ahlering, M. (2020). Evolutionary approaches to seed sourcing for grassland restorations. New Phytologist, 225(6), 2246– 2248. 10.1111/nph.16427

Hapke, A., & Thiele, D. (2016). GIbPSs: a toolkit for fast and accurate analyses of genotyping-by-sequencing data without a reference genome. Molecular Ecology Resources, 16(4), 979– 990. 10.1111/1755-0998.12510

Hausfather, Z., Marvel, K., Schmidt, G. A., Nielsen-Gammon, J. W., & Zelinka, M. (2022). Climate simulations: recognize the ‘hot model’ problem. Nature, 605(7908), 26–29. 10.1038/d41586-022-01192-2

Hausman, C. E., Jaeger, J. F., & Rocha, O. J. (2010). Impacts of the emerald ash borer (EAB) eradication and tree mortality: Potential for a secondary spread of invasive plant species. Biological Invasions, 12(7), 2013–2023. 10.1007/s10530-009-9604-3

Herms, D. A., & McCullough, D. G. (2014). Emerald ash borer invasion of North America: History, biology, ecology, impacts, and management. Annual Review of Entomology, 59, 13–30. 10.1146/annurev-ento-011613-162051

Hijmans, R. J. (2022). geosphere: Spherical Trigonometry. R package version 1.5–18. https://CRAN.R-project.org/package=geosphere

Hoban, S., Bruford, M. W., Funk, W. C., Galbusera, P., Griffith, M. P., Grueber, C. E., Heuertz, M., Hunter, M. E., Hvilsom, C., Stroil, B. K., Kershaw, F., Khoury, C. K., Laikre, L., Lopes-Fernandes, M., MacDonald, A. J., Mergeay, J., Meek, M., Mittan, C., Mukassabi, T. A., … Vernesi, C. (2021). Global commitments to conserving and monitoring genetic diversity are now necessary and feasible. BioScience, 71(9), 964–976. 10.1093/biosci/biab054

Hoban, S., Kelley, J. L., Lotterhos, K. E., Antolin, M. F., Bradburd, G., Lowry, D. B., Poss, M. L., Reed, L. K., Storfer, A., & Whitlock, M. C. (2016). Finding the genomic basis of local adaptation: Pitfalls, practical solutions, and future directions. American Naturalist, 188(4), 379–397. 10.1086/688018

Hohenlohe, P. A., Day, M. D., Amish, S. J., Miller, M. R., Kamps-Hughes, N., Boyer, M. C., Muhlfeld, C. C., Allendorf, F. W., Johnson, E. A., & Luikart, G. (2013). Genomic patterns of introgression in rainbow and westslope cutthroat trout illuminated by overlapping paired-end RAD sequencing. Molecular Ecology, 22(11), 3002–3013. 10.1111/mec.12239

Hollister, J.W. (2023). elevatr: Access Elevation Data from Various APIs. R package version 0.99.0. https://CRAN.R-project.org/package=elevatr/

Hudson, R. R., Slatkin, M., & Maddison, W. P. (1992). Estimation of Levels of Gene Flow From DNA Sequence Data. Genetics, 132, 583–589.

IUCN Standards and Petitions Committee. 2022. Guidelines for Using the IUCN Red List Categories and Criteria. Version 15.1. Prepared by the Standards and Petitions Committee. Downloadable from https://www.iucnredlist.org/documents/RedListGuidelines.pdf

Jaramillo-Correa, J. P., Beaulieu, J., Khasa, D. P., & Bousquet, J. (2009). Inferring the past from the present phylogeographic structure of North American forest trees: Seeing the forest for the genes. Canadian Journal of Forest Research, 39(2), 286–307. 10.1139/X08-181

Jia, K. H., Zhao, W., Maier, P. A., Hu, X. G., Jin, Y., Zhou, S. S., Jiao, S. Q., El-Kassaby, Y. A., Wang, T., Wang, X. R., & Mao, J. F. (2020). Landscape genomics predicts climate change-related genetic offset for the widespread *Platycladus orientalis* (Cupressaceae). Evolutionary Applications, 13(4), 665–676. 10.1111/eva.12891

Jombart, T. (2008). Adegenet: A R package for the multivariate analysis of genetic markers. Bioinformatics, 24(11), 1403–1405. 10.1093/bioinformatics/btn129

Kardos, M., Armstrong, E. E., Fitzpatrick, S. W., Hauser, S., Hedrick, P. W., Miller, J. M., Tallmon, D. A., & Chris Funk, W. (2021). The crucial role of genome-wide genetic variation in conservation. Proceedings of the National Academy of Sciences of the United States of America, 118(48). 10.1073/pnas.2104642118

Kim, M. S., Richardson, B. A., McDonald, G. I., & Klopfenstein, N. B. (2011). Genetic diversity and structure of western white pine (*Pinus monticola*) in North America: A baseline study for conservation, restoration, and addressing impacts of climate change. Tree Genetics and Genomes, 7(1), 11–21. 10.1007/s11295-010-0311-0

Korneliussen, T. S., Albrechtsen, A., & Nielsen, R. (2014). ANGSD: Analysis of Next Generation Sequencing Data. BMC Bioinformatics, 15(1), 1–13.

Korneliussen, T. S., Moltke, I., Albrechtsen, A., & Nielsen, R. (2013). Calculation of Tajima’s D and other neutrality test statistics from low depth next-generation sequencing data. BMC Bioinformatics, 14(1). 10.1186/1471-2105-14-289

Kovacs, K. F., Haight, R. G., McCullough, D. G., Mercader, R. J., Siegert, N. W., & Liebhold, A. M. (2010). Cost of potential emerald ash borer damage in U.S. communities, 2009-2019. Ecological Economics, 69(3), 569–578. 10.1016/j.ecolecon.2009.09.004

Laporte, V., & Charlesworth, B. (2002). Effective population size and population subdivision in demographically structured populations. Genetics, 162(1), 501–519. 10.1093/genetics/162.1.501

Láruson, Á. J., Fitzpatrick, M. C., Keller, S. R., Haller, B. C., & Lotterhos, K. E. (2022). Seeing the forest for the trees: Assessing genetic offset predictions from gradient forest. Evolutionary Applications, 15(3), 403–416. 10.1111/eva.13354

Lasky, J. R., Josephs, E. B., & Morris, G. P. (2023). Genotype-environment associations to reveal the molecular basis of environmental adaptation. Plant Cell, 35(1), 125–138. 10.1093/plcell/koac267

Li, H. (2013). Aligning sequence reads, clone sequences and assembly contigs with BWA-MEM. arXiv:1303.3997

Lind, B. M., Candido-Ribeiro, R., Singh, P., Lu, M., Obreht Vidakovic, D., Booker, T. R., Whitlock, M. C., Yeaman, S., Isabel, N., & Aitken, S. N. (2024). How useful are genomic data for predicting maladaptation to future climate? Global Change Biology, 30(4), 1–19. 10.1111/gcb.17227

Little, E.L., Jr., 1971, Atlas of United States trees: conifers and important hardwoods. 1146, p., 200. U.S. Department of Agriculture.

Liu, J.-J., Sniezko, R., Murray, M., Wang, N., Chen, H., Zamany, A., Sturrock, R. N., Savin, D., & Kegley, A. (2016). Genetic Diversity and Population Structure of Whitebark Pine (*Pinus albicaulis* Engelm.) in Western North America. PLOS ONE, 11(12), e0167986. 10.1371/journal.pone.0167986

Mahony, C. R., MacLachlan, I. R., Lind, B. M., Yoder, J. B., Wang, T., & Aitken, S. N. (2020). Evaluating genomic data for management of local adaptation in a changing climate: A lodgepole pine case study. Evolutionary Applications, 13(1), 116–131. 10.1111/eva.12871

Mahony, C. R., Wang, T., Hamann, A., & Cannon, A. J. (2022). A global climate model ensemble for downscaled monthly climate normals over North America. International Journal of Climatology, 42(11), 5871–5891. 10.1002/joc.7566

McKinney, G. J., Waples, R. K., Pascal, C. E., Seeb, L. W., & Seeb, J. E. (2018). Resolving allele dosage in duplicated loci using genotyping-by-sequencing data: A path forward for population genetic analysis. Molecular Ecology Resources, 18(3), 570–579. 10.1111/1755-0998.12763

McKinney, G. J., Waples, R. K., Seeb, L. W., & Seeb, J. E. (2017). Paralogs are revealed by proportion of heterozygotes and deviations in read ratios in genotyping-by-sequencing data from natural populations. Molecular Ecology Resources, 17(4), 656–669. 10.1111/1755-0998.12613

Mclaughlin, C. M., Hinshaw, C., Sandoval-Arango, S., & Hamilton, J. A. (2024). Redlisting genetics : towards linking genetic data and conservation management. Authorea. April 23, 2024. 10.22541/au.169411992.23557596/v2

Meek, M. H., Beever, E. A., Barbosa, S., Fitzpatrick, S. W., Fletcher, N. K., Mittan-Moreau, C. S., Reid, B. N., Campbell-Staton, S. C., Green, N. F., & Hellmann, J. J. (2023). Understanding Local Adaptation to Prepare Populations for Climate Change. BioScience, 73(1), 36–47. 10.1093/biosci/biac101

Mijangos, J. L., Gruber, B., Berry, O., Pacioni, C., & Georges, A. (2022). dartR v2: An accessible genetic analysis platform for conservation, ecology and agriculture. Methods in Ecology and Evolution, 13(10), 2150–2158. 10.1111/2041-210X.13918

Milligan, B. G. (2003). Maximum-likelihood estimation of relatedness. Genetics, 163(3), 1153– 1167. 10.1093/genetics/163.3.1153

Mimura, M., & Aitken, S. N. (2010). Local adaptation at the range peripheries of Sitka spruce. Journal of Evolutionary Biology, 23(2), 249–258. 10.1111/j.1420-9101.2009.01910.x

Mimura, M., & Aitken, S. N. (2007). Adaptive gradients and isolation-by-distance with postglacial migration in *Picea sitchensis*. Heredity, 99(2), 224–232. 10.1038/sj.hdy.6800987

Munz, P. A., & Laudermilk, J. D. (1949). A Neglected Character in Western Ashes (*Fraxinus*). Aliso: A Journal of Systematic and Floristic Botany, 2(1), 49–62.

Neel, M. C., McKelvey, K., Ryman, N., Lloyd, M. W., Short Bull, R., Allendorf, F. W., Schwartz, M. K., & Waples, R. S. (2013). Estimation of effective population size in continuously distributed populations: There goes the neighborhood. Heredity, 111(3), 189–199. 10.1038/hdy.2013.37

Nei, M., & Li, W. H. (1979). Mathematical model for studying genetic variation in terms of restriction endonucleases. Proceedings of the National Academy of Sciences of the United States of America, 76(10), 5269–5273. 10.1073/pnas.76.10.5269

Nei, M. (1972). Genetic Distance between Populations. The American Naturalist, 106(949), 283–292.

Niemiec, S. S., Ahrens, G. R., Willits, S., & Hibbs, D. E. (1995). Hardwoods of the Pacific Northwest. In Western Hardwoods (Issue March). Forestry Publications Office, Oregon State University.

Nock, C. J., Ovenden, J. R., Butler, G. L., Wooden, I., Moore, A., & Baverstock, P. R. (2011). Population structure, effective population size and adverse effects of stocking in the endangered Australian eastern freshwater cod *Maccullochella ikei*. Journal of Fish Biology, 78(1), 303–321. 10.1111/j.1095-8649.2010.02865.x

Oksanen, J., Simpson, G.L., Blanchet, F. G., Kindt, R., Legendre, P., Minchin, P. R., O’Hara, R. B., Solymos, P., Stevens, M. H. M., Szoecs, E., Wagner, H., Barbour, M., Bedward, M., Bolker, B., Borcard, D., Carvalho, G., Chirico, M., De Caceres, M., Durand, S., Evangelista, H. B. A., FitzJohn, R., Friendly, M., Furneaux, B., Hannigan, G., Hill, M. O., Lahti, L., McGlinn, D., Ouellette, M-H., Cunha, E. R., Smith, T., Stier, A., Ter Braak C. J. F., & Weedon, J. (2022). vegan: Community Ecology Package. R package version 2.6–4. https://CRAN.R-project.org/package=vegan

Oldfield, S. & Westwood, M. 2017. Fraxinus latifolia. The IUCN Red List of Threatened Species 2017: e.T61918519A61918522. 10.2305/IUCN.UK.2017-3.RLTS.T61918519A61918522.en. Accessed on 17 June 2024.

Osuna-Mascaró, C., Agneray, A. C., Galland, L. M., Leger, E. A., & Parchman, T. L. (2023). Fine-scale spatial genetic structure in a locally abundant native bunchgrass (*Achnatherum thurberianum*) including distinct lineages revealed within seed transfer zones. Evolutionary Applications, 16(5), 979–996. 10.1111/eva.13547

Parchman, T. L., Gompert, Z., Mudge, J., Schilkey, F. D., Benkman, C. W., & Buerkle, C. A. (2012). Genome-wide association genetics of an adaptive trait in lodgepole pine. Molecular Ecology, 21(12), 2991–3005. 10.1111/j.1365-294X.2012.05513.x

Pellatt, M. G., Hebda, R. J., & Mathewes, R. W. (2001). High-resolution Holocene vegetation history and climate from Hole 1034B, ODP leg 169s, Saanich Inlet, Canada. Marine Geology, 174(1–4), 211–226. 10.1016/S0025-3227(00)00151-1

Pina-Martins, F., Silva, D. N., Fino, J., & Paulo, O. S. (2017). Structure_threader: An improved method for automation and parallelization of programs structure, fastStructure and MavericK on multicore CPU systems. Molecular Ecology Resources, 17(6), e268–e274. 10.1111/1755-0998.12702

Prive, S. (2016). Overstory Structure and Community Characteristics of Oregon Ash (Fraxinus latifolia) Forests of the Willamette Valley, Oregon. Oregon State University.

Purcell, S., Neale, B., Todd-Brown, K., Thomas, L., Ferreira, M. A. R., Bender, D., Maller, J., Sklar, P., De Bakker, P. I. W., Daly, M. J., & Sham, P. C. (2007). PLINK: A tool set for whole-genome association and population-based linkage analyses. American Journal of Human Genetics, 81(3), 559–575. 10.1086/519795

Puritz, J. B., Hollenbeck, C. M., & Gold, J. R. (2014). dDocent: A RADseq, variant-calling pipeline designed for population genomics of non-model organisms. PeerJ, 2014(1). 10.7717/peerj.431

R Core Team (2024). R: A language and environment for statistical computing. R Foundation for Statistical Computing, Vienna, Austria. https://www.R-project.org/.

Raj, A., Stephens, M., & Pritchard, J. K. (2014). FastSTRUCTURE: Variational inference of population structure in large SNP data sets. Genetics, 197(2), 573–589. 10.1534/genetics.114.164350

Rehfeldt, G. E., Hoff, R. J., & Steinhoff, R. H. (1984). Geographic Patterns of Genetic Variation in *Pinus monticola*. Botanical Gazette, 145(2), 229–239.

Richardson, B. A., Rehfeldt, G. E., & Kim, M. S. (2009). Congruent climate-related genecological responses from molecular markers and quantitative traits for western white pine (*Pinus monticola*). International Journal of Plant Sciences, 170(9), 1120–1131. 10.1086/605870

Ruiz Mondragón, K. Y., Klimova, A., Aguirre-Planter, E., Valiente-Banuet, A., Lira, R., Sanchez-De la Vega, G., & Eguiarte, L. E. (2023). Differences in the genomic diversity, structure, and inbreeding patterns in wild and managed populations of *Agave potatorum* Zucc. used in the production of Tobalá mezcal in Southern Mexico. PLoS ONE, 18(11 November), 1–24. 10.1371/journal.pone.0294534

Savolainen, O., Pyhäjärvi, T., & Knürr, T. (2007). Gene flow and local adaptation in trees. Annual Review of Ecology, Evolution, and Systematics, 38, 595–619. 10.1146/annurev.ecolsys.38.091206.095646

Semizer-Cuming, D., Krutovsky, K. V., Baranchikov, Y. N., Kj r, E. D., & Williams, C. G. (2019). Saving the world’s ash forests calls for international cooperation now. Nature Ecology and Evolution, 3(2), 141–144. 10.1038/s41559-018-0761-6

Soltis, D. E., Gitzendanner, M. A., Strenge, D. D., & Soltis, P. S. (1997). Systematics Chloroplast DNA intraspecific phylogeography of plants from the Pacific Northwest of North America*. Plant Systematics and Evolution, 206, 353–373.

Sniezko, R. A. (2020). Ash (Fraxinus spp.) in western U.S. Summer 2020 Newsletter. https://www.researchgate.net/publication/335755409

St. Clair, J. B., Richardson, B. A., Stevenson-Molnar, N., Howe, G. T., Bower, A. D., Erickson, V. J., Ward, B., Bachelet, D., Kilkenny, F. F., & Wang, T. (2022). Seedlot Selection Tool and Climate-Smart Restoration Tool: Web-based tools for sourcing seed adapted to future climates. Ecosphere, 13(5), 1–18. 10.1002/ecs2.4089

Sun, J., Koski, T. M., Wickham, J. D., Baranchikov, Y. N., & Bushley, K. E. (2024). Emerald Ash Borer Management and Research: Decades of Damage and Still Expanding. Annual Review of Entomology, 69, 239–258. 10.1146/annurev-ento-012323-032231

Sutherland, B. G., Belaj, A., Nier, S., Cottrell, J. E., P Vaughan, S., Hubert, J., & Russell, K. (2010). Molecular biodiversity and population structure in common ash (*Fraxinus excelsior* L.) in Britain: Implications for conservation. Molecular Ecology, 19(11), 2196–2211. 10.1111/j.1365-294X.2009.04376.x

Tabas-Madrid, D., Méndez-Vigo, B., Arteaga, N., Marcer, A., Pascual-Montano, A., Weigel, D., Xavier Picó, F., & Alonso-Blanco, C. (2018). Genome-wide signatures of flowering adaptation to climate temperature: Regional analyses in a highly diverse native range of *Arabidopsis thaliana*. Plant Cell and Environment, 41(8), 1806–1820. 10.1111/pce.13189

Tajima, F. (1989). Statistical method for testing the neutral mutation hypothesis by DNA polymorphism. Genetics, 123(3), 585–595. 10.1093/genetics/123.3.585

Taylor, H. (1945). Cyto-taxonomy and phylogeny of the Oleaceae. Brittonia, 5(4), 337–367.

Twisselmann, E. C. (1967). A flora of Kern county, California. Wasmann Journal of Biology, 25 (1/2), 395.

Vandergast, A. G., Bohonak, A. J., Weissman, D. B., & Fisher, R. N. (2007). Understanding the genetic effects of recent habitat fragmentation in the context of evolutionary history: Phylogeography and landscape genetics of a southern California endemic Jerusalem cricket (Orthoptera: Stenopelmatidae: Stenopelmatus). Molecular Ecology, 16(5), 977–992. 10.1111/j.1365-294X.2006.03216.x

VanWallendael, A., Lowry, D. B., & Hamilton, J. A. (2022). One hundred years into the study of ecotypes, new advances are being made through large-scale field experiments in perennial plant systems. Current Opinion in Plant Biology, 66, 102152. 10.1016/j.pbi.2021.102152

Varas-Myrik, A., Sepúlveda-Espinoza, F., Fajardo, A., Alarcón, D., Toro-Núñez, Ó., Castro-Nallar, E., & Hasbún, R. (2022). Predicting climate change-related genetic offset for the endangered southern South American conifer *Araucaria araucana*. Forest Ecology and Management, 504(October 2021). 10.1016/j.foreco.2021.119856

Wang, I. J., & Bradburd, G. S. (2014). Isolation by environment. Molecular Ecology, 23(23), 5649–5662. 10.1111/mec.12938

Wang, T., Hamann, A., Spittlehouse, D., & Carroll, C. (2016). Locally downscaled and spatially customizable climate data for historical and future periods for North America. PLoS ONE, 11(6), 1–17. 10.1371/journal.pone.0156720

Waples, R. K., Larson, W. A., & Waples, R. S. (2016). Estimating contemporary effective population size in non-model species using linkage disequilibrium across thousands of loci. Heredity, 117(4), 233–240. 10.1038/hdy.2016.60

Waples, R. S. (2022). What Is N_e_, Anyway? Journal of Heredity, 113(May), 371–379.

Waples, R. S. (2024). Practical application of the linkage disequilibrium method for estimating contemporary effective population size: A review. Molecular Ecology Resources, 24(1), 1–16. 10.1111/1755-0998.13879

Waples, R. S. (2010). Spatial-temporal stratifications in natural populations and how they affect understanding and estimation of effective population size. Molecular Ecology Resources, 10(5), 785–796. 10.1111/j.1755-0998.2010.02876.x

Waples, R. S., Antao, T., & Luikart, G. (2014). Effects of overlapping generations on linkage disequilibrium estimates of effective population size. Genetics, 197(2), 769–780. 10.1534/genetics.114.164822

Waples, R. S., & Yokota, M. (2007). Temporal estimates of effective population size in species with overlapping generations. Genetics, 175(1), 219–233. 10.1534/genetics.106.065300

Watterson, G. A. (1975). On the Number of Segregating Sites in Genetical Models without Recombination. Theoretical Population Biology, 276(7), 256–276.

Zheng, X., Levine, D., Shen, J., Gogarten, S. M., Laurie, C., & Weir, B. S. (2012). A high-performance computing toolset for relatedness and principal component analysis of SNP data. Bioinformatics, 28(24), 3326–3328. 10.1093/bioinformatics/bts606

