## Supplemental for "Genomics-driven monitoring of *Fraxinus latifolia* (Oregon Ash) for conservation and EAB-resistance breeding"

**Supplemental Methods**

In the southern portion of the *Fraxinus latifolia* distribution, hybridization with *F. velutina* Torr. has been reported and may involve tetraploid populations (Taylor, 1945; Munz and Laudermilk, 1949; Twisselmann, 1967). Thus, the potential for polyploidy was evaluated using the ‘gbs2ploidy’ v1.0 R package (Gompert & Mock, 2017) in R version 4.4.0 (R Core Team, 2024). Posterior probabilities of allelic proportions were estimated using the *estprops* function based on two expected ploidy levels (diploid and tetraploid) with 10,000 MCMC steps, a 1000 step burn-in, an MCMC thinning of 100. The *estploidy* function was then used to classify samples in groups representing the putative ploidy levels. The *estploidy* function performs a principal component analysis (PCA) on the estimated allelic proportions from *estprops* followed by a discriminant analysis to identify clusters of samples in principal component space and classify them into groups representing potential ploidy levels, providing probability values of ploidy level for each sample.

**Supplemental Results**

To explore factors that may contribute to the distinct genetic clusters across the species distribution (Fig. 1; Fig. S1) we used a PCA of allelic proportions to estimate ploidy level variation among individuals (Fig. S2). The majority of populations clustered together, while populations representing the southern Sierra Nevada mountains and southern disjunct distribution comprised a distinct, but smaller cluster with PC2 (26.1%) largely describing the distinction between northern and southern populations’ allelic proportions. Predicted ploidy level based on *estploidy* suggested tetraploids persist in eight southern populations, with six populations entirely tetraploid (AKR, KRL, MRY, SGR, SIE, and SNF of CA), and two others that represent a mix between primarily diploid individuals and several putative tetraploids (HUM of California, and WPD of Oregon). Ploidy level grouping probabilities (ie., predicted probability of a sample being either diploid or tetraploid) were extremely high for members of each respective cluster (>0.99), except for one sample of the WPD, OR population (group 1, diploid: 0.187, group 2, tetraploid: 0.813; Table S2).

**Supplemental Literature Cited**

Gompert, Z., & Mock, K. E. (2017). Detection of individual ploidy levels with genotyping-by-sequencing (GBS) analysis. *Molecular Ecology Resources*, *17*(6), 1156–1167. https://doi.org/10.1111/1755-0998.12657

Munz, P. A., & Laudermilk, J. D. (1949*). A Neglected Character in Western Ashes* (*Fraxinus*). ). *Aliso: A Journal of Systematic and Floristic Botany, 2*(1), 49–62.

Taylor, H. (1945). Cyto-taxonomy and phylogeny of the Oleaceae. *Brittonia*, *5*(4), 337–367.

Twisselmann, E. C. (1967). A flora of Kern county, California. *Wasmann Journal of Biology*, 25 (1/2), p. 395.

R Core Team (2024). R: A language and environment for statistical computing. R Foundation for Statistical Computing, Vienna, Austria. URL https://www.R-project.org/.

**Figure S1.** Map (A) and visualization of principal components 2 and 3 for all populations (B) and the subset of populations used in all analyses (C).

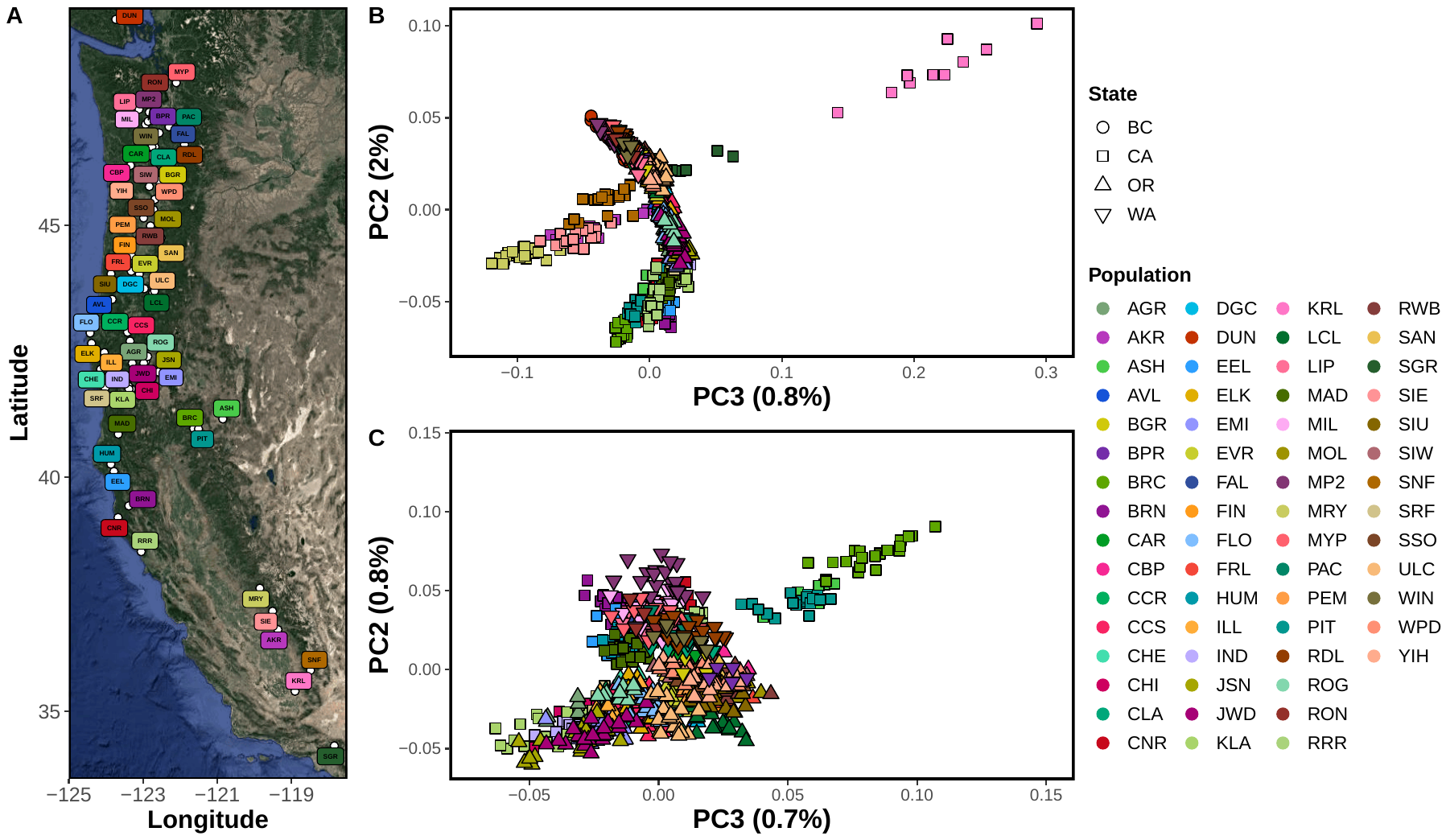

**Figure S2.** Results from PCA on allelic proportions used to estimate ploidy level. Two clusters are identified within the plot - putative tetraploid populations from the southern extent of the range and two northern samples and a putative diploid cluster comprising samples from all other populations. Group probabilities were generally very high (>0.95), except for one sample designated to the putative tetraploid group (WPD, OR; group 2 probability = 0.813).

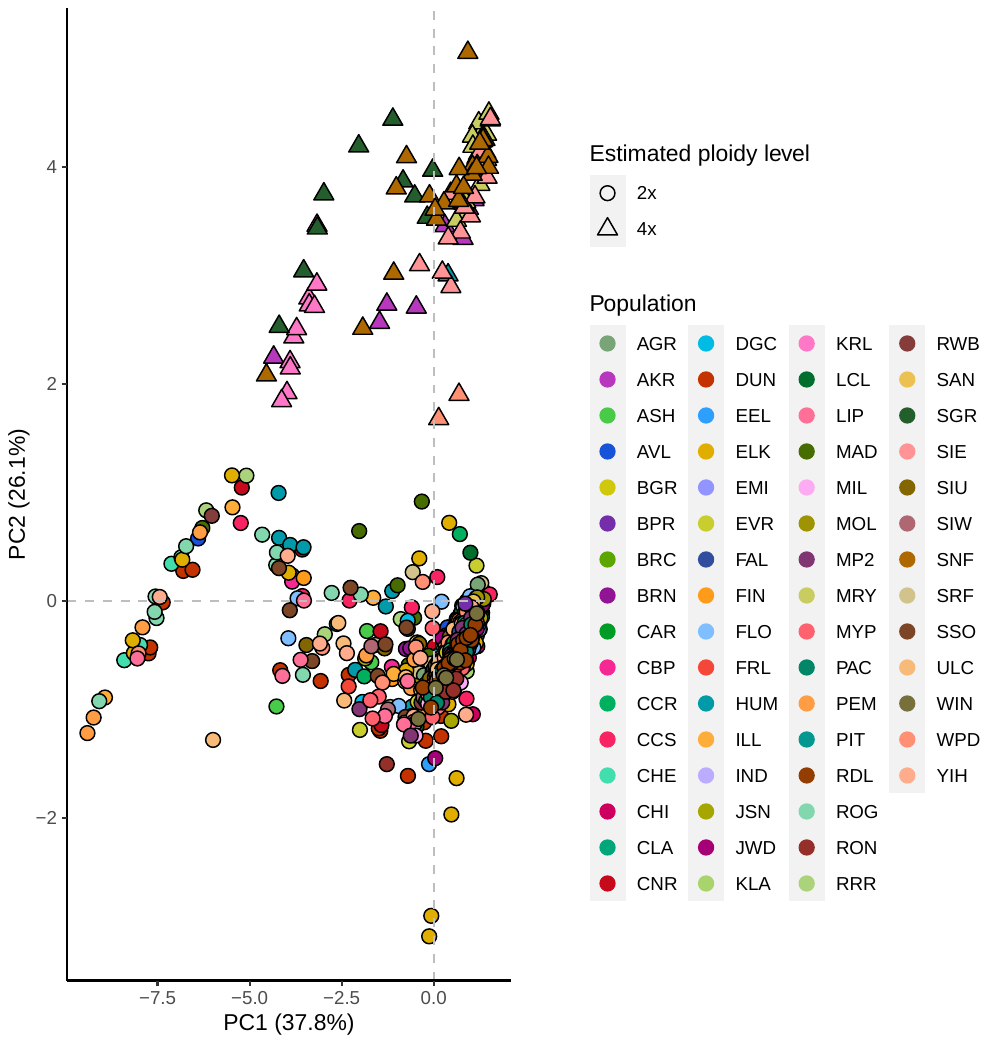

**Figure S3.** Visualization of ancestry Q-scores from *structure* analysis (A) and marginal likelihood scores for *k*’s two through six (B). A north-to-south pattern of clustering is identified for all clusters, similar to PCA results. In *k* = 4, the British Columbia, Canada population forms a distinct cluster (dark red), all Washington, USA and northern Oregon, USA populations from the Columbia River watershed form a cluster (yellow), southern Oregon and northern California populations from the Klamath-Siskiyou Ecoregion form a cluster (red), and all other California populations form a cluster (blue). A *k* of two had the lowest marginal likelihood score and a *k* of four had the highest (B).

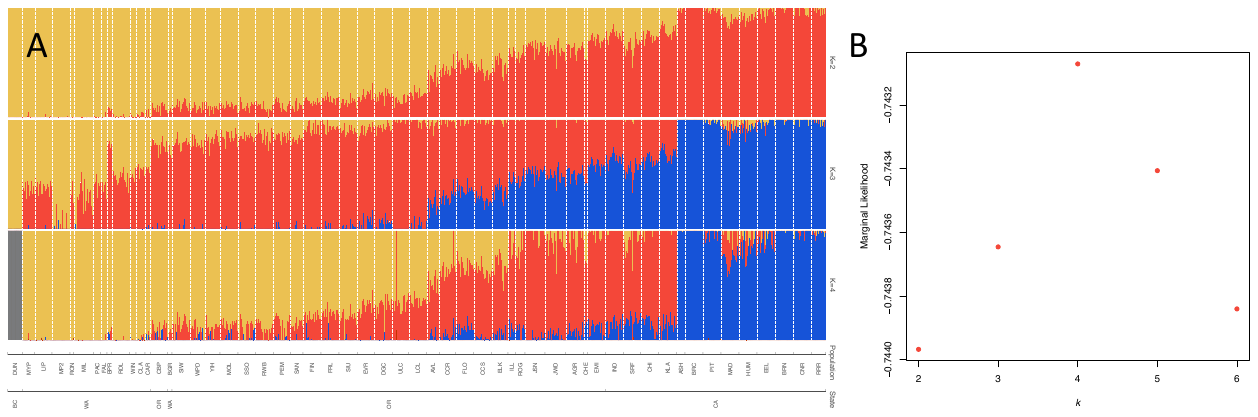

**Figure S4.** Nucleotide diversity (π), Watterson’s Θ (ΘW), and Tajima’s *D* (D). Nucleotide diversity (π; A) and (ΘW; B) were generally very low across populations ranging from 0.0034 (π) and 0.0023 (ΘW) to 0.0055 (π) and 0.0051 (ΘW). Tajima’s *D* values were negative for all populations but one and are consistent with recent population expansion after a genetic bottleneck.

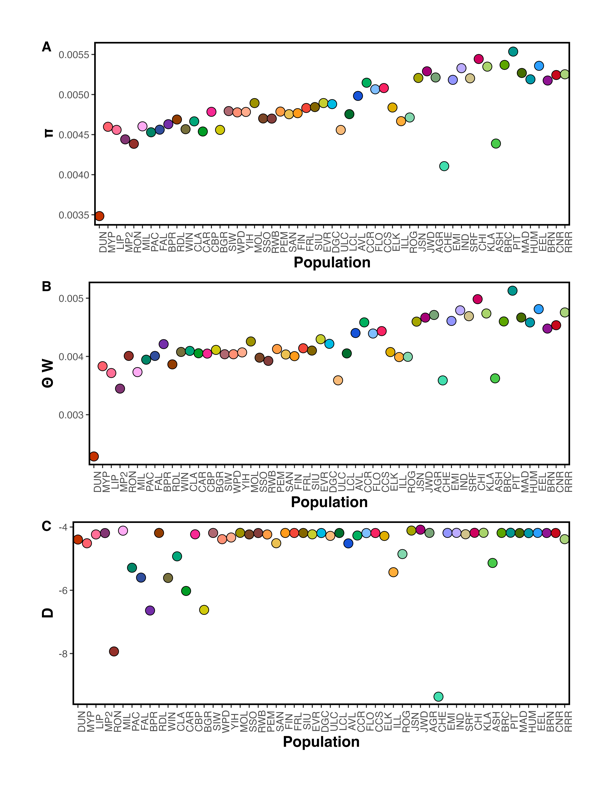

**Figure S5.** Heatmap of *F*_ST_ (upper triangle) and Nei’s *D* (lower triangle) values. *F*_ST_ values ranged from 0.030 (CCR, Oregon, USA vs. CCS, Oregon, USA) to 0.242 (DUN, British Columbia, Canada vs. CHE, Oregon, USA). Nei’s *D* ranged from 0.010 (CCR, Oregon, USA vs. CCS, Oregon, USA) to 0.094 (CHE, Oregon, USA vs. DUN, British Columbia, Canada).

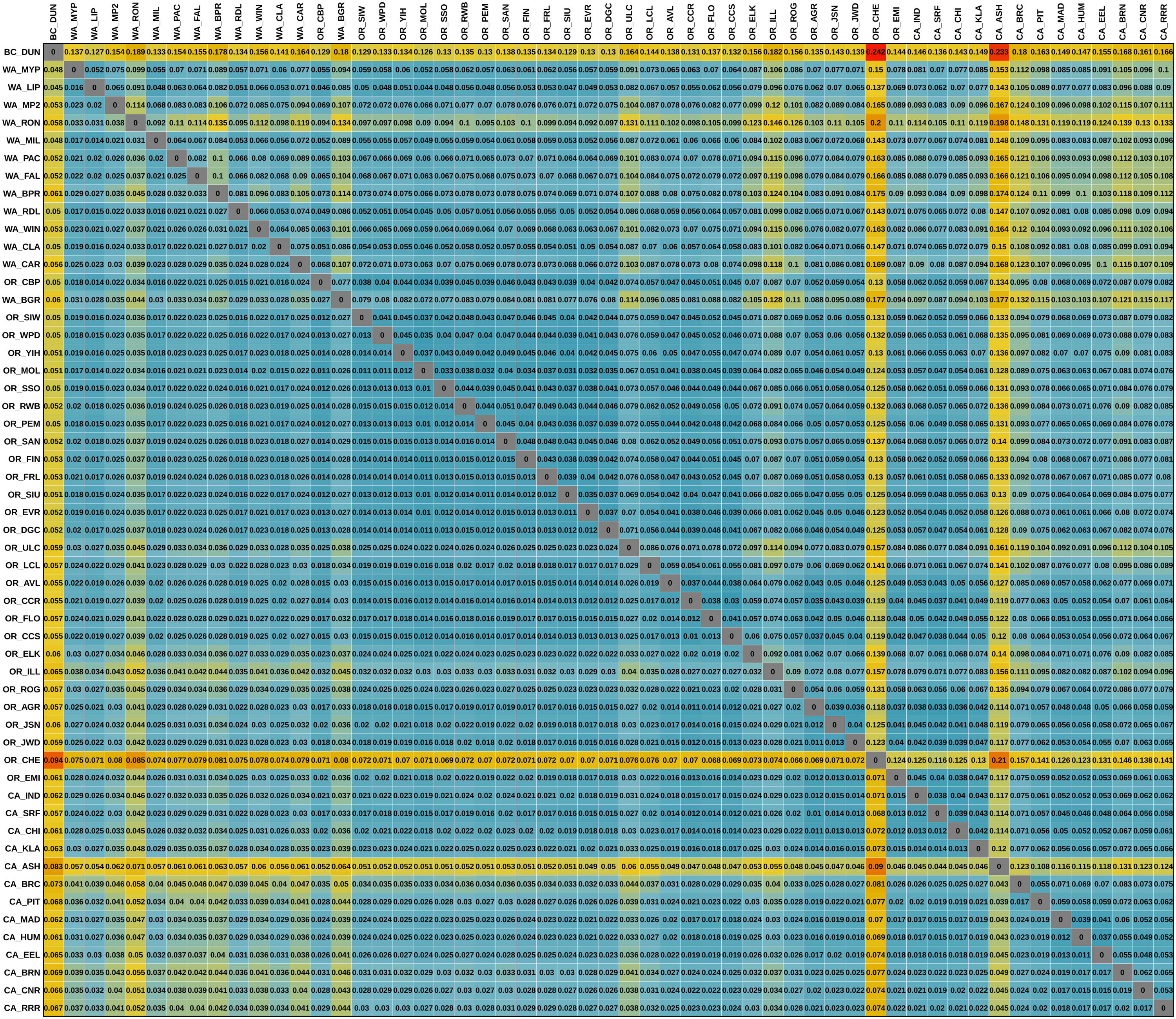

**Figure S6.** Observed heterozygosity (*H*_O_) is predicted to be greatest along the major river system valleys central to the species distribution. *H_O_* ranged from 0.143±0.189 (CHE, Oregon, USA) to 0.244±0.165 (CHI, California, USA) with a mean of 0.226±0.014.

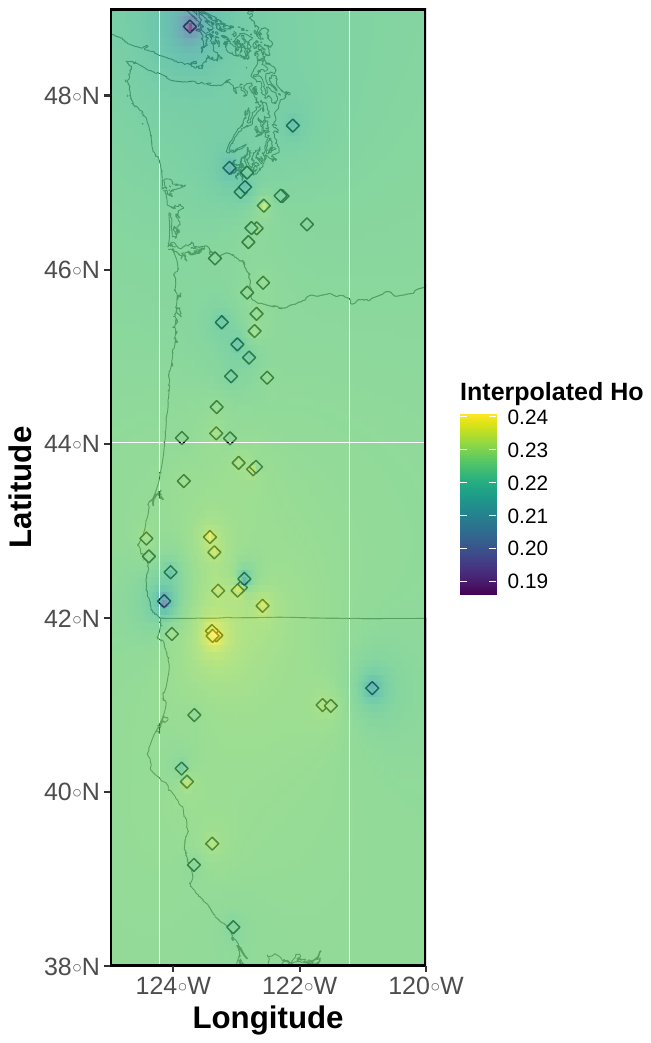

**Figure S7.** Predicted genetic offsets were consistent across all shared socio-economic pathways (SSP) and ranged from 0.0869 to 0.130 for an optimistic climate change scenario (ssp245; A) and 0.0869 to 0.134 for an extreme climate change scenario (ssp585; C).

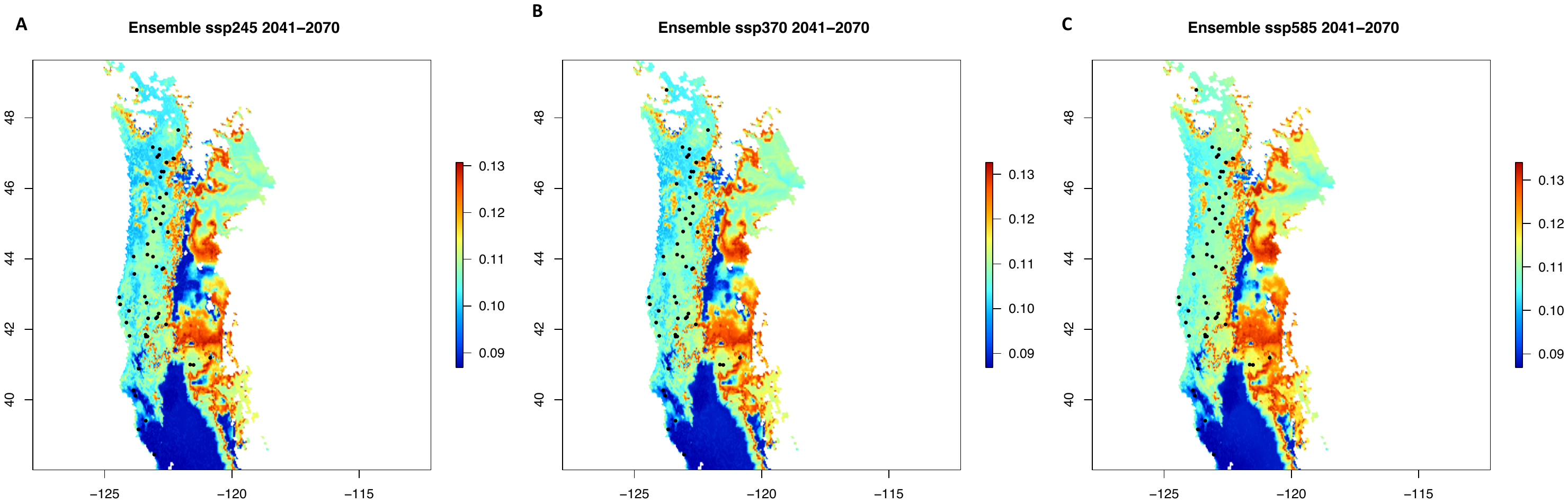

**Figure S8.** (A) Percent of samples with known sex per population, with the mean percent indicated by dashed grey line (69.02%). (B) Number of female and male samples per population.

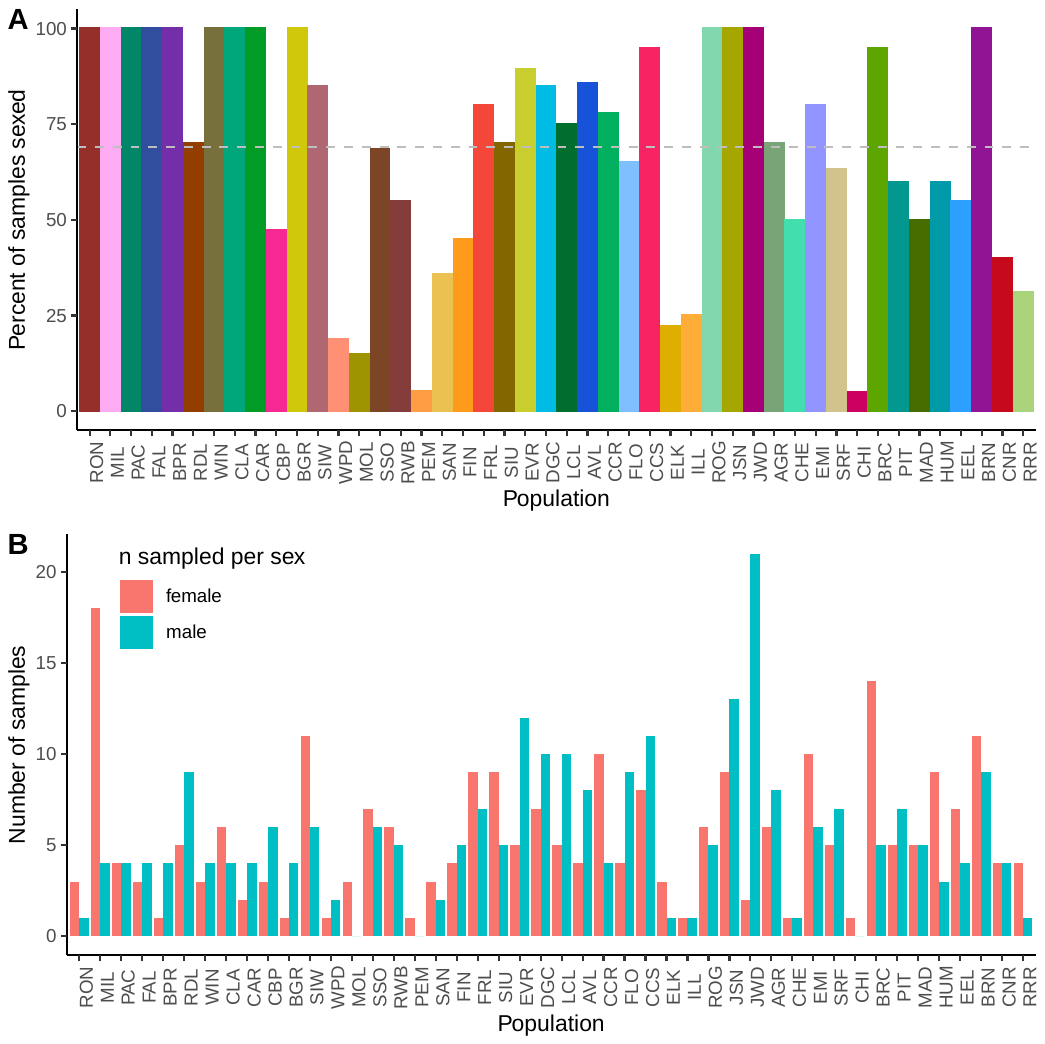

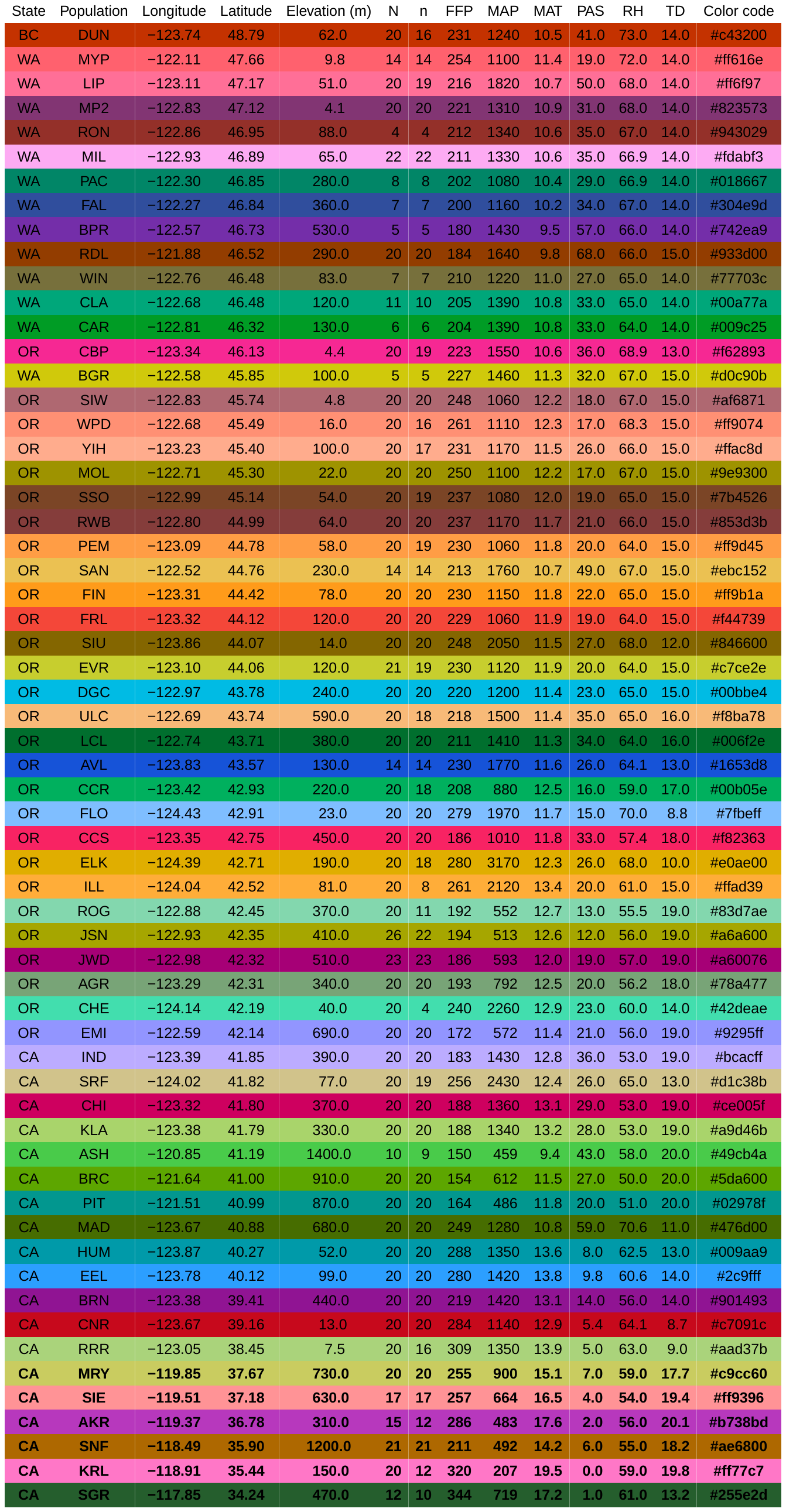

Table S1**.** Final sampling data for each population that was successfully sequenced and passed quality control, ordered by decreasing latitude. Six populations of the southern terminus of the range were excluded post- SNP and allelic frequency PCA. Latitude and longitude represent an average for all samples within the population. Environmental variables represent all uncorrelated variables used in analyses: Frost-free period (FFP; days), mean annual precipitation (MAP; mm), mean annual temperature (MAT; °C), precipitation as snow (PAS; mm), relative humidity (RH; %), and continentality (TD; °C).
